# Host and microbiome jointly contribute to adaptation to a complex environment

**DOI:** 10.1101/2023.03.22.533853

**Authors:** Carola Petersen, Inga K. Hamerich, Karen L. Adair, Hanne Griem-Krey, Montserrat Torres Oliva, Marc P. Hoeppner, Brendan J.M. Bohannan, Hinrich Schulenburg

## Abstract

Most animals and plants have associated microorganisms, collectively referred to as their microbiomes, which can provide essential functions. Given their importance, host-associated microbiomes have the potential to contribute substantially to adaptation of the host-microbiome assemblage (the “metaorganism”). Microbiomes may be especially important for rapid adaptation to novel environments because microbiomes can change more rapidly than host genomes. However, it is not well understood how hosts and microbiomes jointly contribute to metaorganism adaptation. We developed a model system with which to disentangle the contributions of hosts and microbiomes to metaorganism adaptation. We established replicate mesocosms containing the nematode *Caenorhabditis elegans* co-cultured with microorganisms in a novel complex environment (laboratory compost). After approximately 30 nematode generations (100 days), we harvested worm populations and associated microbiomes, and subjected them to a common garden experiment designed to unravel the impacts of microbiome composition and host genetics on metaorganism adaptation. We observed that adaptation took different trajectories in different mesocosm replicates, with some increasing in fitness and others decreasing, and that interactions between host and microbiome played an important role in these contrasting evolutionary paths. We chose two exemplary mesocosms (one with a fitness increase and one with a decrease) for detailed study. For each example, we identified specific changes in both microbiome composition (for both bacteria and fungi) and nematode gene expression associated with each change in fitness. Our study provides experimental evidence that adaptation to a novel environment can be jointly influenced by host and microbiome.

## Introduction

Most animals and plants have associated microorganisms, collectively referred to as their microbiomes, which can provide important biological functions. These functions include digestion of otherwise indigestible materials (e.g. (1)), production of essential nutrients (e.g. (2)), increased resistance to pathogens (e.g. (2–4)), and stimulation of development (including maturation of the immune system; e.g. (5)), among many others. Because of the potential impact of microbiomes on crucial physiological functions, it has been suggested that multicellular organisms are best conceptualized as “metaorganisms” – multispecies assemblages with collective properties such as fitness (6).

Given the importance of microbiomes to metaorganism function, they could play an important role in evolutionary adaptation, especially in response to environmental change. Microbiomes may be especially important in mediating acclimation and adaptation to environmental change because microbiomes can rapidly respond to environmental challenges, both through changes in microbiome composition and through genetic and phenotypically plastic changes in individual microbial lineages. Hosts may respond more slowly than their microbiomes to a changing environment because hosts often have longer generation times and smaller population sizes than their associated microorganisms (7).

Microbiome-mediated acclimation to environmental change has been documented in multiple metaorganisms, most frequently in response to increasing temperatures (8). For example, some corals have been found to harbor heat-tolerant microbes, which can help the coral survive in warmer waters, as reported for *Acropora hyacinthus* (9). Similarly, long-term exposure of the sea anemone *Nematostella vectensis* to increased temperatures led to both higher heat tolerance and microbiome changes; subsequent transplant experiments demonstrated that the higher heat tolerance was a consequence of changes in microbiome composition (10).

Even if more slowly, genetic changes in the host population (i.e., host evolution) could still improve performance of the metaorganism in the new environment. To date, a joint assessment of the contribution of either microbiome acclimation and/or host evolution has only rarely been attempted. One of the few examples is the study of pathogen stress in the nematode host *Caenorhabditis elegans. C. elegans*, together with a single symbiont *Enterococcus faecalis*, was exposed to pathogen stress over 14 host generations under controlled laboratory conditions. The symbiont was observed to evolve an increased protective effect and simultaneously the host evolved an increased ability to accommodate the protective symbiont (11–13). A more recent example adapted the parasitoid wasp *Nasonia vitripennis* with its diverse microbiome over 85 generations to the herbicide atrazine, demonstrating that both changes in the microbiome as well as genetic adaptations in the host increased atrazine resistance in an interdependent manner, consistent with a co-adapted host-microbiome association (14). In this example, it is as yet unclear whether the genetic changes in the host favor colonization with the beneficial microbes and/or directly mediate resistance (14). Overall, the microbiome can play a central role in metaorganism acclimation to novel environmental conditions, yet to date the importance of host genetic adaptation in this context is poorly understood.

The aim of our study is to establish a novel experimental metaorganism system for studying the causes and consequences of microbiome-mediated acclimation, and to specifically explore the contribution of both host and microbiome to improved performance in a novel environment. We developed and implemented a mesocosm experimental approach with the nematode *C. elegans* and its microbiome as a model. This nematode is common in temperate regions across the world, where it proliferates in rotting plant matter, especially rotting fruits or compost (15), where it associates with a species-rich gut microbiome, consisting of Proteobacteria such as Enterobacteriaceae and *Pseudomonas, Stenotrophomonas, Ochrobactrum*, and *Sphingomonas* bacteria, as well as certain yeast species (16,17). We maintained a genetically diverse *C. elegans* laboratory population in an experimental compost environment similar in many respects to the nematode’s natural habitat (15,18). After 100 days (approx. 30 host generations), we harvested and separated worm populations and associated microbiomes, and subjected them to a common garden experiment designed to unravel the impacts of microbiome composition and host genetics on metaorganism adaptation. We observed that adaptation took different trajectories in different mesocosm replicates, with some increasing in fitness and others decreasing, and that interactions between host and microbiome played an important role in these contrasting evolutionary paths.

## Materials and Methods

### Nematode and bacterial strains

The mesocosm experiment was initiated with the experimental and genetically diverse, androdioecious *C. elegans* population A_0_ derived from 16 inter-crossed natural isolates (19). We used a set of 43 bacterial strains, labeled CeMbio43, as an initial inoculum for the mesocosm experiment and as reference standard for the common garden experiments. The CeMbio43 community consists of bacterial strains, which were isolated from natural *C. elegans* and its substrate and which are representative of the native *C. elegans* microbiome (see full list in Supplementary Table S1 (16,17,20,21). The CeMbio43 community is identical to the one used by us in a separate study, with the exception of three strains that have been added to the mixture in the separate work (i.e., strains *Stenotrophomonas* sp. MYb526 and MYb536, and *Chryseobacterium* sp. MYb317).

### Mesocosm experiment

To assess adaptation of the metaorganism, we set up a mesocosm experiment, in which the genetically diverse A_0_ *C. elegans* population and the initial inoculum of the CeMbio43 bacteria were subjected over 100 days to a non-sterile environment consisting of decomposing fruits and vegetables (Figure 1A; see supplement for more details). This compost environment has not been experienced by the experimental A_0_ population, yet it is related to the natural habitat of *C. elegans* (15,18), thereby representing a generally suitable context for the worms. See supplementary information for details on preparation of laboratory compost and collection of worms and bacteria.

**Figure 1:**
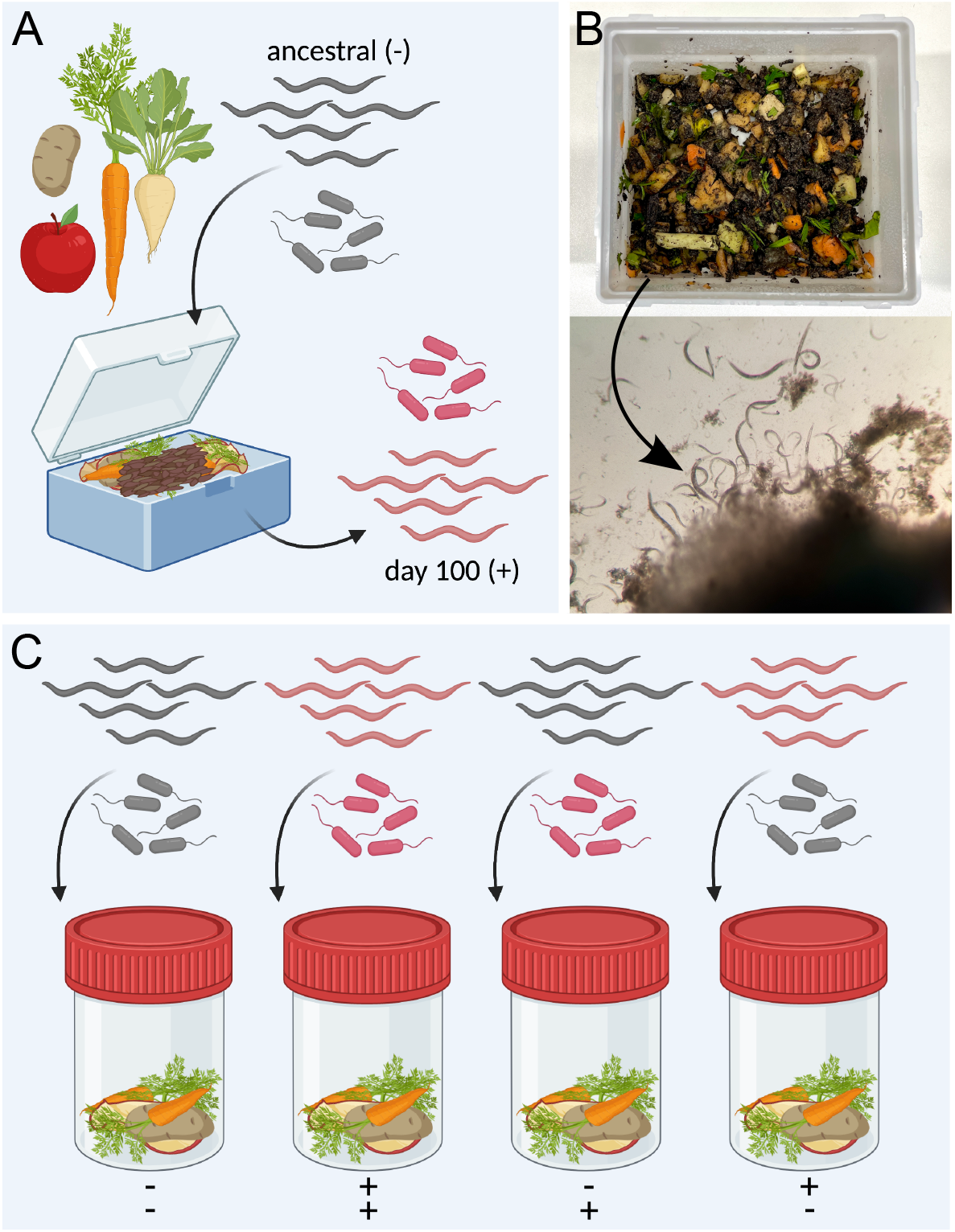
Mesocosm and common garden experiments. **(A)** For the mesocosm experiment, a genetically diverse, ancestral *C. elegans* population (−) and a reference microbiome (−) were allowed to adapt to a laboratory compost environment. *C. elegans* populations (+) and microbial communities (+) from six mesocosms were isolated at day 100. **(B)** The initial compost of the mesocosm experiment consisted of compost soil and plant material (top). Worms from proliferating mesocosm populations were isolated from compost samples covered with a buffer (bottom). **(C)** In a common garden experiment, the ancestral *C. elegans* population and the initial reference microbiome were combined with the day-100 nematode populations and the corresponding day-100 microbial communities (from the same mesocosm replicate) in all possible four combinations in either a compost environment (as illustrated) or an agar plate (not shown). **(A, C)** Created with BioRender.com.

### Common garden experiment and assessment of nematode population growth rate

To determine how the host-microbiome assemblage adapted to the new compost environment, we isolated nematode populations and microbial communities from the six mesocosm replicates at day 100 and tested them in a common garden experiment. We performed two sets of common garden experiments: the first with material from all mesocosm replicates (using only 1 technical replicate), and the second with only material from replicates Box 1 and Box 2 (using five replicates per treatment combination). For each common garden experiment, we combined the ancestral A_0_ nematode population and the initial microbial inoculum as reference with the day-100 nematode populations and the corresponding day-100 microbial community (from the same mesocosm replicate) in all possible four combinations in either a compost (both sets of common garden experiments) or on agar plates (only the second), the latter used as an alternative environment known to be suitable for nematode proliferation from the multitude of *C. elegans* studies. Each experiment was followed by an assessment of *C. elegans* population growth (both common garden experiments) and also nematode length and nematode area (only the second experiment) as proxies for host fitness. For the second set of common garden experiments, we additionally used the obtained material for an analysis of microbial community composition using both 16S and ITS amplicon sequence analysis (for bacteria and fungi, respectively) and also an analysis of the *C. elegans* transcriptome response. See supplementary information for details on preparation of laboratory compost, collection of worms and bacteria, and statistics.

### 16S and ITS amplicon sequencing for microbiome analysis of common garden experiment

Microbial community composition was characterized for substrate samples and nematodes, collected at the end of the second set of compost common garden experiments, involving Box 1 and Box 2. After DNA isolation, we used 16S rRNA gene and ITS amplicon sequencing to determine the relative abundance of bacteria and fungi, respectively, following established protocols for sequencing, data processing, and statistical analyses, as outlined in detail in the supplementary information.

### RNAseq for transcriptome analysis of *C. elegans* populations

We assessed the transcriptomic response of *C. elegans* populations from all treatment combinations of the second set of common garden experiments with compost. Compost and worms for transcriptomics were prepared separately but using the same general approach as for the population growth assay in five replicates. Worms were isolated after 24 h, followed by RNA isolation, RNAseq and transcriptome data analyses, following established protocols (22,23), as outlined in detail in the supplementary information.

## Results

### Novel compost mesocosm supports stable proliferating *C. elegans* populations

We developed a novel protocol for the long-term maintenance of proliferating populations of *C. elegans* in laboratory compost mesocosms. We exposed a genetically diverse *C. elegans* population to a reference microbiome in laboratory mesocosms consisting of decomposing plant material (i.e. non-sterile chopped vegetables) and soil (Figure 1A, B; Supplementary Figure S1). Regular addition of plant material served to supply the mesocosm microbiomes with nutrients and in turn led to consistently high worm counts. Using this protocol, we maintained proliferating worm populations (Figure 1B; Supplementary Movies 1-3) for more than 500 days (equivalent to approx. 150 *C. elegans* generations; mesocosm experiment still ongoing), demonstrating that the mesocosm compost provides suitable conditions for stable growth of *C. elegans* under semi-natural conditions.

### Nematode fitness in the compost environment is influenced by both host and microbiome

To determine how the nematode and microbial populations responded to the compost environment, we focused on an analysis of worms and microbiomes harvested after 100 days (equivalent to approx. 30 *C. elegans* generations; Figure 1B). We co-inoculated different combinations of nematodes and microbiomes into common environments (“common garden” experiments; Figure 1C). At the conclusion of these experiments, we measured several components of nematode fitness, including population growth rate and nematode size (i.e. length and area; see Methods above).

Our first common garden experiment consisted of inoculating day-100 mesocosm worm populations or ancestral worm populations into fresh compost, either with the corresponding day-100 microbiome (from the same mesocosm replicate) or with the reference microbiome used for the initial inoculation of the mesocosms. Overall, measures of worm fitness did not differ between day-100 worms and ancestral worms (Figure 2A; Supplementary Table 1). However, we observed substantial differences among the independent mesocosm replicates, particularly when combined with different microbiomes. We chose two exemplary mesocosm replicates with contrasting patterns, labeled Box 1 and Box 2, for subsequent analyses (Figure 2A). During this first common garden experiment, the Box 1 day-100 worms produced high numbers of offspring, especially when inoculated with their respective day-100 microbiomes, whereas ancestral worms produced almost no offspring with the same microbial inoculum. In contrast, Box 2 day-100 worms produced few offspring when inoculated with their day-100 microbiomes or with the reference microbiome, whereas ancestral worms produced many offspring when inoculated with these same microbiomes. These results suggest that the relative contribution of host and microbiome to metaorganism fitness may have diverged significantly between Box 1 and Box 2 over the 100 days of this experiment. However, this conclusion is based on single replicates of each combination of worm population and microbiome.

**Figure 2.**
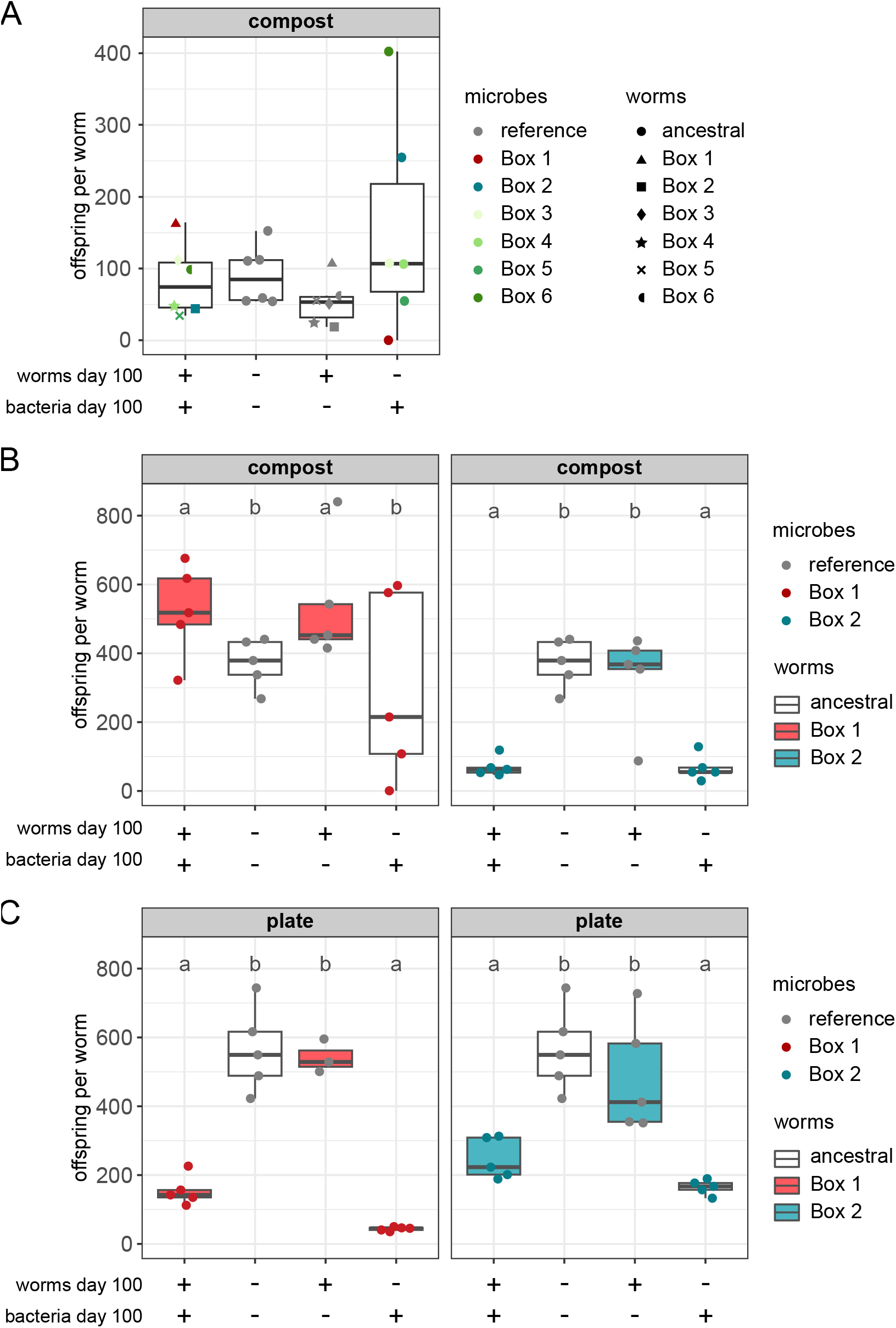
Host and microbiome can jointly determine nematode fitness in the novel compost environment. Results of common garden experiments, in which population growth was measured for *C. elegans* populations isolated from mesocosms at day 100 (+) and ancestral worms (−) in the presence of mesocosm day-100 microbiomes (+) or the reference microbiomes including the CeMbio43 bacterial community (−). Population growth is shown as offspring per worm. **(A)** Population growth of six mesocosm replicates (Boxes 1-6) and one ancestral worm population, measured under compost conditions. Colors indicate different mesocosm microbiomes and the reference microbiomes (gray); symbols indicate worm populations from the different mesocosm boxes and the ancestral worm population. n = 1. **(B)** Population growth of Box 1 day-100 (red boxes), Box 2 day-100 (blue boxes), and ancestral worm populations (white boxes) in the presence of Box 1 day-100 (red dots) or Box 2 day-100 (blue dots) microbiomes or reference microbiomes (gray dots). n = 5 **(C)** Population growth of Box 1 day-100 (red boxes), Box 2 day-100 (blue boxes) and ancestral worms (white boxes) on agar plates with Box 1 day-100 (red dots), Box 2 day-100 (blue dots) or reference microbiomes (gray dots). Results are summarized as boxplots with the median as a thick horizontal line, the interquartile range as box, the whiskers as vertical lines, and each replicate depicted by a dot or symbol. n = 5.

In the second set of common garden experiments we asked whether this conclusion was robust to replication. These experiments focused only on the Box 1 and Box 2 worm populations and microbiomes (excluding those from the other mesocosms). We assessed the fitness of the day-100 worm populations and the ancestral worm population, each combined with either day-100 microbiomes or the reference microbiome. We conducted these experiments in a compost environment (as before) and additionally on agar plates, in order to determine whether changes in fitness were specific to the compost environment.

In Box 1 compost, the number of worm offspring produced was significantly influenced by the nature of the worm population (i.e. day-100 vs ancestral; *p* = 0.027; Supplementary Table S1), whereas the size of individual worms (i.e. worm area) was affected by the type of microbial inoculum (i.e. day-100 vs reference microbiomes; *p* = 0.015; Supplementary Figure S2; Supplementary Table S1). As in the first common garden experiment, the Box 1 day-100 worms produced many offspring when combined with their day-100 microbiome (Figure 2B). The number of offspring of the ancestral worms inoculated with the Box 1 day-100 microbiome varied substantially among replicates (Figure 2B). In contrast, the Box 2 day-100 microbial inoculum consistently and significantly reduced worm offspring numbers, worm length, and worm area produced by both the Box 2 day-100 and the ancestral worms (in all cases, *p* < 0.01; Figure 2B; Supplementary Table S1). These results confirm that the Box 1 nematode population has increased in fitness following 100 days in the laboratory compost environment, while the Box 2 population has decreased in fitness. Furthermore, these results suggest that these fitness differences are at least in part due to differences in the Box 1 and Box 2 microbiomes.

On agar plates, the microbial inoculum significantly influenced the offspring numbers, worm length and worm area (*p* < 0.01; Figure 2C; Supplementary Figure S2; Supplementary Table S1). For Box 1 worms, the results on agar differed from those in compost; offspring numbers were higher on plates containing the reference microbiome than those with Box 1 day-100 microbiomes, and ancestral worms produced the fewest offspring when inoculated with Box 1 day-100 microbiomes (Figure 2C). For Box-2 worm populations, the number of offspring per worm was significantly influenced by the type of the microbial inoculum (*p* < 0.001; Supplementary Table S1), similar to the results observed in compost (Figure 2B, C). These results suggest that at least the changes in Box 1 metaorganism fitness observed after 100 days in laboratory compost were specific to the compost environment.

### Microbiome treatments resulted in differences in both compost and nematode microbiomes

To determine whether the mesocosm replicates Box 1 and Box 2 varied in microbiome composition, we focused on the common garden experiments in which ancestral worms were exposed to three distinct microbiome treatments (i.e. Box 1 microbiomes, Box 2 microbiomes or the reference microbiomes) in the compost environment. Microbiome treatment was the strongest influence on microbiome composition, across both worm and substrate samples, explaining over 30% of the variation among these samples for both the bacteria and fungi (Figure 3A; Supplementary Table S2). Worm and substrate samples also differed from each other in microbiome composition; however, this depended on the mesocosm replicate. Specifically, worm and substrate microbiomes differed significantly when ancestral worms were exposed to the reference microbiome (16S, *R*^*2*^ = 0.25, *p* = 0.03; ITS, *R*^*2*^ = 0.29, *p* = 0.03) or Box 2 microbiomes (16S, *R*^*2*^ = 0.42, *p* = 0.04; ITS, *R*^*2*^ = 0.27, *p* = 0.03) but not the Box 1 microbiomes (16S, *R*^*2*^ = 0.18, *p* = 0.14; ITS, *R*^*2*^ = 0.11, *p* = 0.74). These results suggest that the substrate microbiome from the Box 1 treatment contains microbes that are able to colonize *C. elegans* and/or are preferentially taken up by the nematodes. In contrast, the Box 2 microbiome treatment results in significantly different substrate and worm microbiomes, indicating the presence of microbes that cannot colonize nematodes and/or are avoided by *C. elegans*.

**Figure 3.**
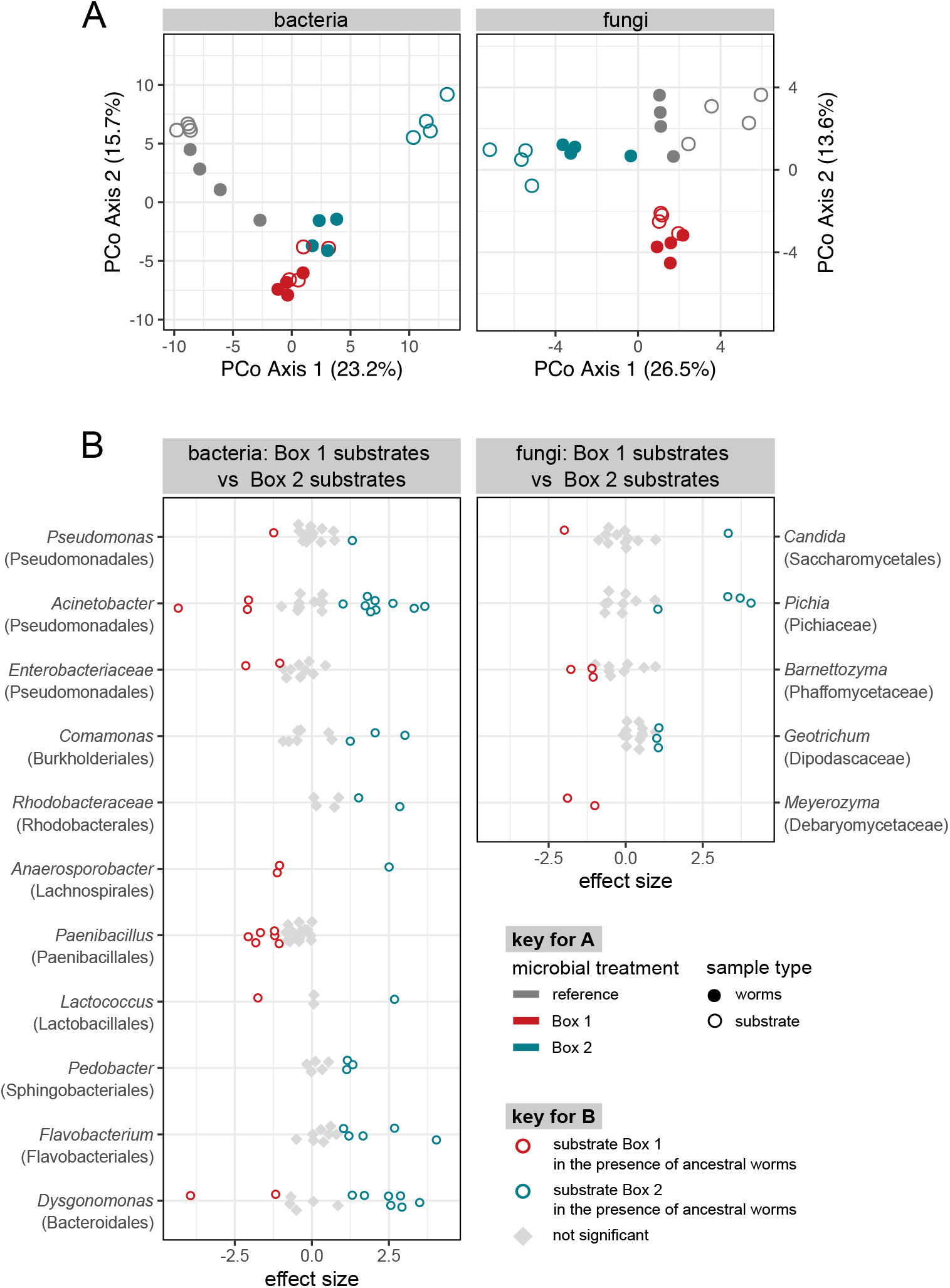
Microbiome treatments resulted in differences in both compost and nematode microbiomes. **(A)** Ordination (Principal Coordinates analysis of Aitchison distance) depicting variation in microbiome composition across inoculation treatments in a common garden experiment. **(B)** Differential abundance analysis of substrate microbiomes inoculated with Box 1 day-100 microbiome or Box 2 day-100 microbiome.

We next investigated the specific taxonomic differences between the Box 1 and Box 2 substrate microbiomes and identified both bacterial and fungal taxa (i.e., ASVs) with differential abundance (Figure 3B). For example, ASVs from the fungal genera *Pichia* and *Geotrichum* and the bacterial genera *Comamonas, Pedobacter*, and *Flavobacterium* were more abundant in substrates receiving the Box 2 inoculum. Substrates receiving the Box 1 inoculum were enriched in ASVs from the fungal genera *Barnettozyma* and *Meyerozyma* and the bacterial genera *Paenibacillus* and *Anaerosporobacter* (Figure 3B). A subset of these taxa was differentially abundant between ancestral worms exposed to the Box 1 inoculum and those exposed to the Box 2 inoculum (Supplementary Figures S3, S4).

### Microbiome community changes are associated with differences in nematode fitness

We next focused on understanding the microbiome contributions to the increased fitness observed for Box 1 day-100 worms. To accomplish this, we compared treatments with either ancestral worms or Box 1 day-100 worms in combination with either the reference microbiome inoculum or the Box 1 day-100 inoculum. Across these treatments microbiome composition differed significantly between inocula type for both the substrate and worms. Worm genotype did not influence the microbiome composition of either worms or substrate samples (Figure 4A; Supplementary Table S2). Microbiome composition was very similar between worm and substrate samples in those treatments that received the Box 1 day-100 inoculum. Worm and substrate microbiomes were also similar in those treatments that received the reference inoculum, but to a lesser degree. This pattern was consistent for both fungal and bacteria microbiomes (Figure 4A; Supplementary Table S2). To identify microbes that may contribute to the higher fitness observed for the Box 1 day-100 worms exposed to Box 1 day-100 microbiome, we looked for ASVs that were differentially abundant in Box 1 day-100 worms exposed to either the co-existing Box 1 day-100 microbiome or the reference microbiome. Those worms exposed to the Box 1 day-100 microbiome had a consistently higher relative abundance of ASVs from the fungal genus *Barnettozyma* and bacterial genera *Paenibacillus* and *Dysgonomonas*, and a lower relative abundance of *Sphingobacterium* (Figure 4B). Some bacterial genera (e.g., *Pseudomonas* and *Acinetobacter*) exhibited an inconsistent response, with some ASVs within the genus being more and others less abundant in worms with the Box 1 day-100 inoculum. This result suggests that differences among microbial species or even strains may have important consequences for the increased fitness we observed. When worms exposed to the Box 1 day-100 inoculum were compared to their respective substrates, we observed few differentially abundant ASVs and no consistent taxonomic differences between the sample types (Supplementary Figures S5, S6). We next turned our attention to identifying microbiome contributions to the decreased worm fitness associated with the Box 2 microbiome. To address this, we compared treatments with either ancestral worms or worms from the Box 2 mesocosms in combination with the reference inoculum or microorganisms from the Box 2 mesocosms. Microbiome composition differed significantly between inoculum types for worm and substrate samples and there was no influence of worm genotype on microbiome composition, similar to what was observed in the analysis of the Box 1 treatment combinations. However, while worm and substrate samples tended to be similar (i.e. cluster together; Figure 4A) for the Box 1 treatment combinations, we observed significant separation of worm and substrate microbiomes when exposed to the microbes from the Box 2 mesocosms (Figure 5A). To determine which microbial taxa underlie this separation, we compared microbiomes of worms and substrate samples from the Box 2 microbiome treatments. Relative to worms, substrates had higher relative abundance of ASVs from the bacterial genera *Sphingobacterium, Flavobacterium*, and *Dysgonomonas*. ASVs from the bacterial genera *Pectobacterium, Anaerosporobacter*, and *Enterococcus* and the fungal genera *Pichia* had a higher relative abundance in the worm samples relative to the substrate samples (Figure 5B). Most of these same enriched taxa were observed regardless of whether ancestral or Box 2 day-100 worms were compared to substrate (Supplementary Figures S7, S8).

**Figure 4.**
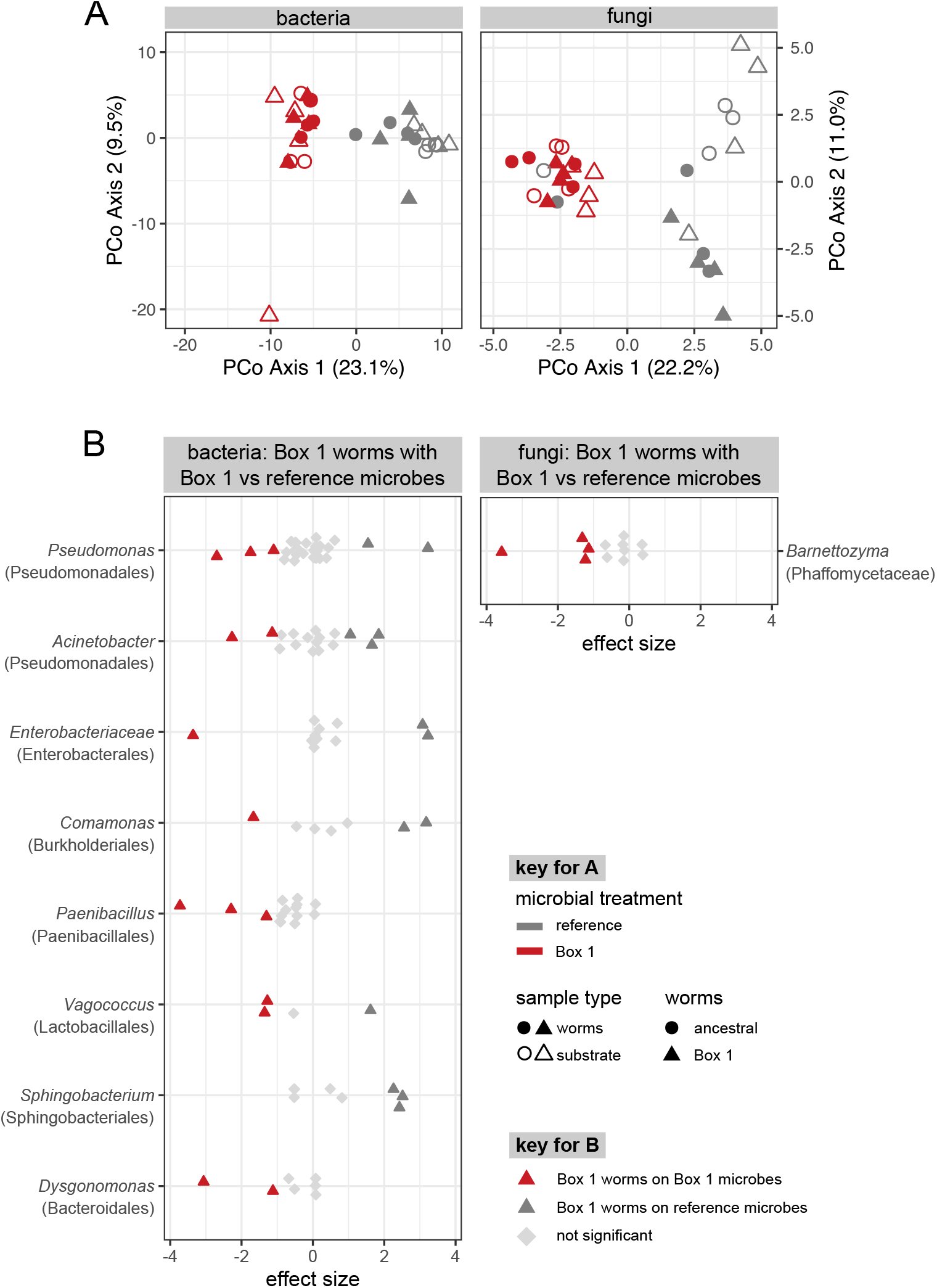
Differences in microbiome composition were associated with increased fitness in nematodes from the Box 1 mesocosm. **(A)** Ordination (Principal Coordinates analysis of Aitchison distance) depicting variation in microbiome composition across treatments in a common garden experiment. **(B)** Differential abundance analysis of microbiomes from Box 1 day-100 worms exposed to the Box 1 day-100 microbiome or the reference microbiome.

**Figure 5.**
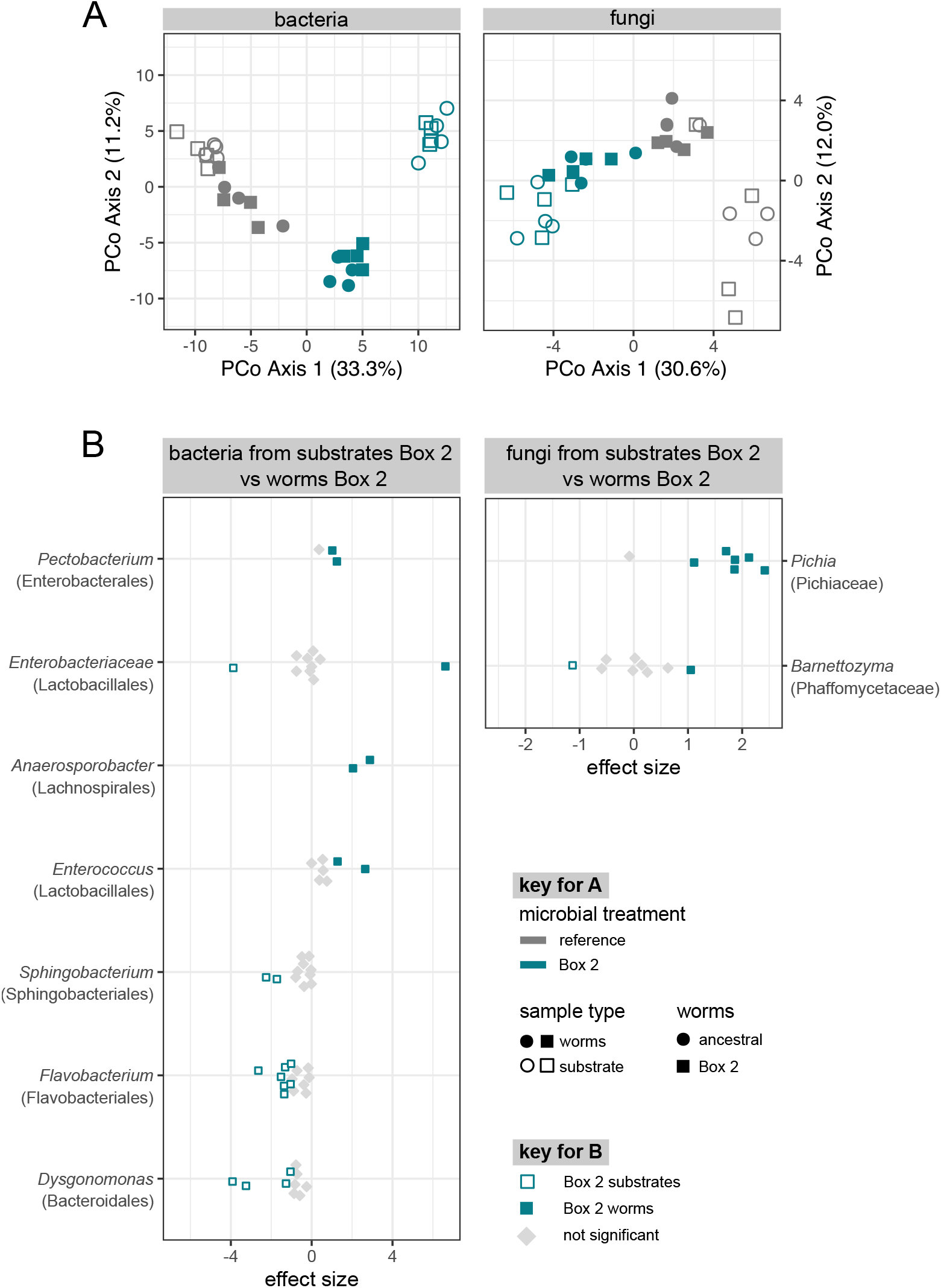
Differences in microbiome composition were associated with decreased fitness in nematodes from the Box 2 mesocosm. **(A)** Ordination (Principal Coordinates analysis of Aitchison distance) depicting variation in microbiome composition across treatments in a common garden experiment. **(B)** Differential abundance analysis of substrate and worm microbiomes exposed to the Box 2 day-100 microbiome.

### Gene expression differs between different nematode populations inoculated with the same microbiomes

It is clear from our common garden experiments that the Box 1 nematode population has increased in fitness following 100 days in the laboratory compost environment, while the Box 2 population has decreased in fitness. It is also clear that these differences in fitness are associated with differences in microbiome composition. To determine whether these changes in fitness were also associated with changes in the host population, we compared variation in gene expression among the Box 1, Box 2 and ancestral *C. elegans* populations after exposure to the reference microbiome in the compost environment. Assessing gene expression in a common environment and after exposure to identical microbiomes allowed us to isolate host genetic changes from those induced by environmental differences.

An explorative PCA revealed a clear separation of the Box 1 and Box 2 worm populations along the first principal component (PC1, explaining 27.8% of the variation), whereas the ancestral population diverged from the two others along PC2 (21.1% variation; Figure 6A; Supplementary Figure S9B). A subsequent *k-means* clustering of the significantly differentially expressed genes yielded four distinct clusters, of which clusters 2 and 4 indicated genes that show contrasting expression patterns for Box 1 and Box 2 nematode populations (Figure 6B; Supplementary Table S3). Cluster 4 consisted of a single gene, *srh-178*, encoding a transmembrane G protein-coupled receptor belonging to the class H serpentine receptors with no specific known function, that is upregulated in the Box 2 but downregulated in the Box 1 worms. Cluster 2 included 41 genes, which we used for a focused enrichment analysis. While the DAVID analysis did not indicate any clearly enriched categories (Supplementary Figure S9A), the *C. elegans*-tailored gene expression analysis with WormExp showed an overrepresentation of gene sets with few main functions, including genes known to be differentially regulated among different natural *C. elegans* strains and additionally stress response as well as longevity genes, which are generally downregulated in the Box 2 but not Box 1 worms (Figure 6C; Supplementary Figure S9C; Supplementary Table S3). These results strongly suggest that the two *C. elegans* populations did indeed change genetically and thus evolved in their mesocosms. Moreover, they diverged from each other, whereby the Box 2 worms showed a reduction in stress response and lifespan genes under these standardized conditions. We next asked whether each of the separate common garden experiments indicates a host transcriptome response that is consistent with the observed phenotypic variation.

**Figure 6.**
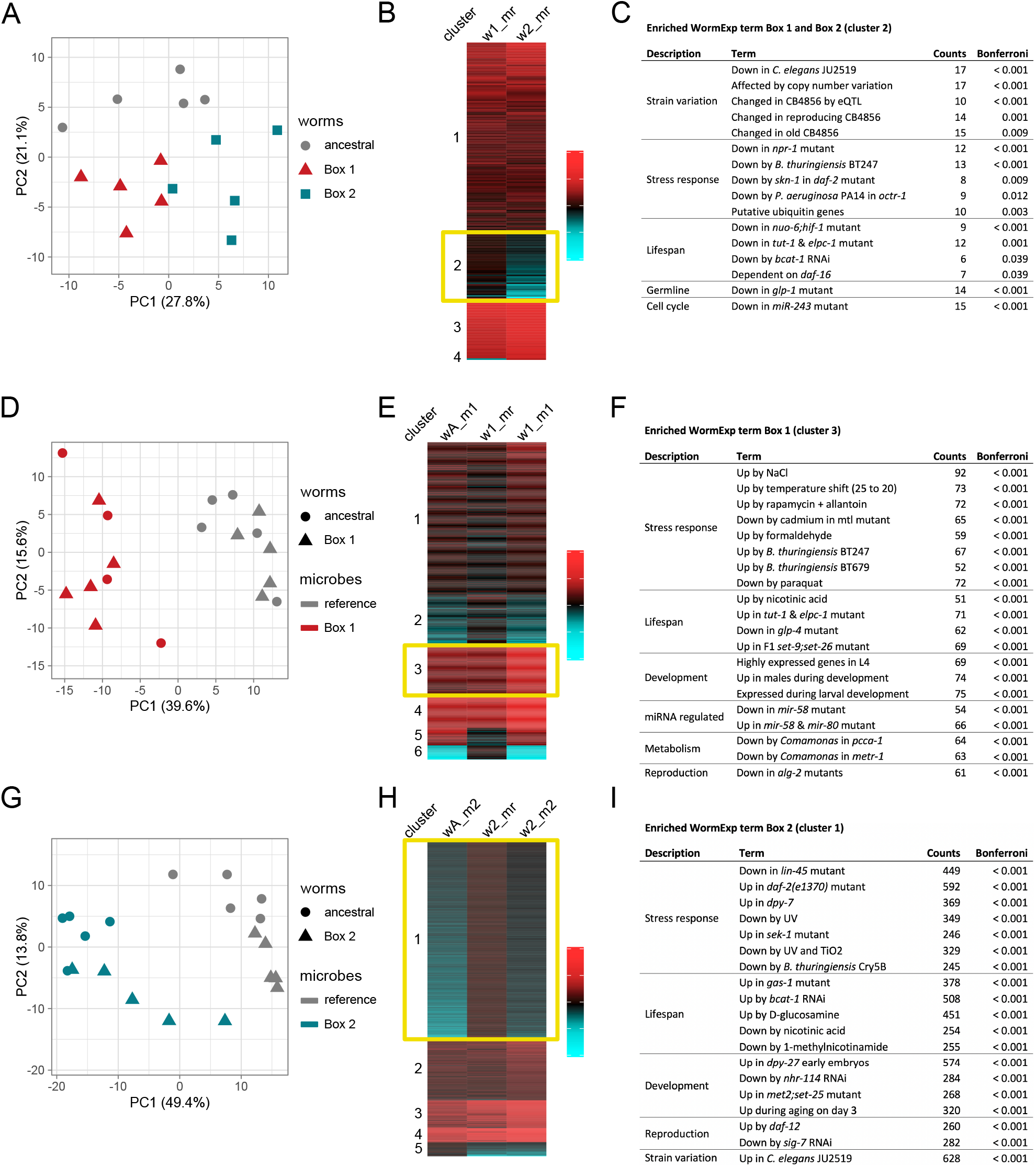
Differential gene expression in the adapted Box 1 and Box 2 *C. elegans* populations. Transcriptome data analysis for the three main sets of analyses, including **(A-C)** a comparison of ancestral, Box 1 day-100 and Box 2 day-100 *C. elegans* populations assayed under identical compost conditions with the reference microbiomes including the CeMbio43 bacterial community, **(D-F)** a comparison of all possible host-microbiome combinations for the Box 1 common garden experiment, and **(G-I)** a comparison of all possible host-microbiome combinations for the Box 2 common garden experiment. In the latter two cases, ancestral or Box 1/Box 2 day-100 worms were combined with either the reference microbiomes or the Box 1/Box 2 day-100 microbiomes. In all three cases, general variation in gene expression was explored with a principal component analysis, whereby panels **(A, D, G)** show the spread of sample variation along the first two principal components (PC1, PC2). Thereafter, variation in significantly differentially expressed genes was assessed using *k*-means clustering and visualization of differential expression using heatmaps, whereby the heatmaps always show fold change of gene expression relative to the ancestral worms combined with reference microbiomes **(B, E, H)**. The clusters, which were used for a focused enrichment analysis, are highlighted by a yellow rectangle. Abbreviations: wA, ancestral worm population; w1, Box 1 day-100 worm population; w2, Box 2 day-100 worm population; mr, reference microbiome; m1, Box 1 day-100 microbiome, m2, Box 2 day-100 microbiome. **(C, F, I)** show the results of the enrichment analysis with the *C. elegans*-tailored WormExp database. Description in the first column indicates overarching functions for the enriched gene sets, for which terms are given in the second column. The third column shows the number of genes of the query set that are responsible for the enrichment with the indicated gene set, while the fourth column shows the inferred probability of enrichment, after Bonferroni correction.

### Gene expression differs between identical nematode populations inoculated with different microbiomes

To determine whether microbiome composition influenced host gene expression, we compared variation in gene expression among the Box 1, Box 2 and ancestral *C. elegans* populations after exposure to different microbiomes in the compost environment. For Box 1, we compared the gene expression of the day-100 nematode population or the ancestral population inoculated with either the Box 1 day-100 microbiome or the reference microbiome in a full factorial design. The initial explorative PCA indicated a clear separation by microbiome type along PC1 (Figure 6D, explaining 36.9% variation). Worm populations (Box 1 day-100 vs ancestral) were separated along PC3 (Supplementary Figure 10B, explaining 9.7% variation). Subsequent *k*-means clustering identified six distinct clusters (Figure 6E). Of these, cluster 3 produced the most convincing pattern of a distinct expression profile for Box 1 worms exposed to Box 1 microbiomes in comparison to all others, in this case consisting of a pronounced upregulation of 88 genes. To specifically explore which functions account for the high performance of the Box 1 worms with Box 1 microbiomes, we focused the enrichment analyses on only this cluster 3. The DAVID analysis revealed an enrichment of the GO terms pseudopodium, carbohydrate binding, cytoskeleton, and cytoplasm, all with medium importance (Supplementary Figure 10A). The WormExp analysis further identified an enrichment of numerous gene sets, including the strong upregulation of genes involved in distinct stress responses (different stress response genes and categories than above), lifespan extension, and development (Figure 6F; Supplementary Figure 10C; Supplementary Table S3). These results strongly indicate that the high fitness of Box 1 worms colonized with Box 1 microbes in the compost environment is associated with a pronounced stress response that targets distinct abiotic as well as biotic stresses (e.g., osmotic stress, heavy metals, toxic substances, and pathogens, Figure 6F; Supplementary Table S3). In addition, high fitness under these conditions is associated with upregulation of longevity and developmental genes.

For Box 2, we again used a full factorial design and compared the gene expression of either the day-100 or ancestral nematode population inoculated with either the Box 2 day-100 microbiome or the reference microbiome. We observed that inoculation with the Box 2 microbiome resulted in the downregulation of distinct stress responses in both the Box 2 as well as the ancestral *C. elegans* populations. Exposure to the Box 2 microbiomes produces a clearly distinct gene expression in both host populations, as revealed by the explorative PCA (Figure 6G; Supplementary Figure 11B) and also the differential gene expression analysis, indicated especially by cluster 1 of the *k*-means clustering analysis (Figure 6H). This cluster 1 includes downregulation of 782 genes. Using DAVID, cluster 1 is enriched for a large number of GO terms, especially those for regulation of transcription and DNA binding (Supplementary Figure 11A). The WormExp analysis similarly revealed enrichment of numerous gene sets (Figure 6I; Supplementary Figure 11C), ultimately indicating the downregulation of genes involved in distinct stress responses (e.g., UV and pathogen stress), lifespan, development, and reproduction. Overall, these results suggest that the microbial community from Box 2 compromises nematode stress responses, most likely explaining the observed decrease in worm fitness, irrespective of *C. elegans* population.

## Discussion

We established a novel experimental metaorganism model, which enables the long-term cultivation of *C. elegans* populations and the assessment of metaorganism adaptation to a complex environment. To date, *C. elegans* has been used for numerous experimental evolution studies (24), focused for example on the assessment of mutation accumulation under relaxed selection (25), adaptation to fluctuating environments (26), or host-pathogen coevolution (27,28). All of these studies have been performed under highly artificial laboratory conditions, where worms are maintained on agar plates or in a standardized liquid medium, usually supplemented with a single food bacterium, *Escherichia coli* OP50, which it does not encounter in nature (15). In all of these studies, *C. elegans* is maintained without its microbiome, which is removed via a bleaching protocol that kills bacteria but not nematode eggs. The model we describe here permits analysis of

*C. elegans* with its microbiome in a structured compost environment similar to its natural habitat (15,18). A compost environment was used previously for *C. elegans*-microbiome studies, but only for short-term experiments (29). In our model, the compost environment was inoculated with a reference microbiome including the CeMbio43 bacterial community, a representative mixture of bacteria from the native microbiome of *C. elegans* (Supplementary Table 1; (17,21)). Since the mesocosm was by design maintained under non-sterile conditions, additional microorganisms colonized the mesocosms in addition to this mixture. This set-up allowed us to explore how the *C. elegans* metaorganism adapted to the compost environment.

We observed that adaptation took different trajectories in different mesocosm replicates over 100 days, with some increasing in fitness and others decreasing. This is remarkable, given that all of the replicate mesocosms were initiated with the same worm population, the same initial bacterial inoculum and the same starting plant material, and were maintained under identical conditions. We do not know in detail what caused these different trajectories, but they could be driven by random mutation events in the host genome, stochastic microbiome assembly across the mesocosm replicates, and/or subtle differences among the replicates in starting conditions, among many possible causes.

We identified one exemplary test case for further study, mesocosm Box 1, which strongly increased in fitness (Figure 2B). We identified genetic changes in the *C. elegans* population (Figures 6A, 6B) and compositional changes in the associated microbiome (Figures 3, 4) that coincided with the increase in fitness. The host evolved by increasing its general stress response (Figures 6C, 6F), possibly allowing it to cope well with a complex compost environment that is much more structurally and physiologically challenging than the agar plate environment it previously experienced. The results from the common garden experiment suggest that these host genetic changes lead to increased fitness irrespective of the microbiome inoculum (Figure 2B). Nevertheless, the coexisting microbiome from the Box 1 mesocosm has a highly specific influence on fitness of the adapted worm population from Box 1, because it only causes an increase in nematode population growth in the compost environment but not on agar plates, leading to the highest median fitness values in compost (Figures 2B, 2C). Thus, our study provides one of the first experimental demonstrations of a joint host genetic and microbiome effect on metaorganism adaptation. Most previous work focused on the contribution of the microbiome or host genome alone to adaptation (discussed in (7,30)), with the exception of the recent study in *Nasonia* wasps, for which metaorganism adaptation to the herbicide atrazine was associated with changes in both host genetics and microbiome composition (14).

We identified both bacterial and fungal lineages associated with this increase in fitness, including lineages that contain microbial species known to be beneficial to *C. elegans*; for example, bacteria from the genus *Pseudomonas* isolated from natural populations of *C. elegans* have been shown to have beneficial effects (16,31,32). There may have been specific evolutionary adaptations in individual microbial lineages that resulted in their positive impact on the fitness of the worm populations that evolved in Box 1; however, this remains a topic for future research since we cannot determine this solely from our amplicon data.

Although we do not yet know exactly how these microbial taxa from Box 1 influence worm fitness, we do have evidence that the Box 1 microbiome as a whole is well-adapted to association with the *C. elegans* population that evolved in the Box 1 mesocosm, because the compositions of the substrate microbiomes and worm microbiomes resulting from inoculation with microbiomes from the Box 1 mesocosm are very similar to each other (Figure 3A). This is not the case for other worm and inoculum combinations, which result in worm and substrate microbiomes that are less similar or in some cases substantially different (Figure 3A). This pattern is consistently present for both fungi and bacteria.

Adaptation through changes to both the microbiome and host are only one of the outcomes that we observed in our experimental system. We also observed a maladaptive response (i.e. a decrease in metaorganism fitness). We chose one mesocosm that exemplifies this response, Box 2, for further study. The worms from this replicate exhibited decreased fitness in the compost environment, but only when inoculated with the substrate microbiome from the Box 2 mesocosm. This inoculant caused substantially decreased fitness in the ancestral worm population as well. It also resulted in a fitness reduction in the agar plate environment for both the ancestral and the Box 2 worm population, although not to the same degree as in compost. These results suggest that the microbiome that developed in this mesocosm was generally detrimental to worm health.

We identified both bacterial and fungal lineages associated with this detrimental effect on fitness, which included members of the bacterial genera *Flavobacterium* and *Dysgonomonas*, and the fungal genus *Pichia*. Although we do not yet know how these microbes decrease metaorganism fitness, we do have evidence that the Box 2 microbiome is not well-adapted overall to association with *C. elegans*, because the compositions of the worm microbiomes and substrate microbiomes resulting from inoculation with microbiomes from the Box 2 mesocosm are very different from each other (Figure 3A). This is true for both fungi and bacteria. This is in contrast to the situation that results from inoculation with the Box 1 microbiome, as described above. This pattern is consistent with the hypothesis that the Box 2 microbiome contains members that are poorly adapted to association with the worm host.

Using our novel *C. elegans*/compost system, we have established that changes in both host and microbiome can jointly mediate metaorganism adaptation. Our experimental system is sufficiently tractable that it will be possible, in future work, to quantify the relative contributions of both host and microbiome to the adaptive process. Adaptive evolution is often based on quantitative genetic changes, yet to date, it is unknown how precisely the associated microbiome interacts with host genetics to determine fitness. A theoretical framework of an extended quantitative genetics model including a term for the microbiome has been published (33). However, it has not yet been applied, most likely for practical reasons, as it requires experimental assessment of defined host lineages with known genetic relationship in experimentally controlled association with microbial isolates (33). Our *C. elegans* model is especially well suited for such analyses, considering that the nematode can be easily crossed (with its generation time of approx. 3 days), and then combined with isolates of its associated, usually aerobic microbes under highly controlled conditions. Such a dissection of the adaptive process may further allow us to reconstruct whether the evolutionary changes in the host directly target the new environmental challenge (e.g., via increased expression of protective molecules) or rather increase the uptake and maintenance of beneficial microbes under the new conditions, equivalent of a microbiome-mediated Baldwin effect (30).

## Acknowledgements

We thank the Bohannan and Schulenburg groups for discussions and advice on this work. For sequencing and amplicon analysis, we further thank the Competence Centre for Genomic Analysis (CCGA) Kiel (funded via the DFG Research Infrastructure NGS_CC project 407495230 as part of the Next Generation Sequencing Competence Network project 423957469), especially Corinna Bang, Sören Franzenburg and the members of project Z3 of CRC 1182. The *C. elegans* A_0_ population was originally provided by Henrique Teotónio, Paris, France. For the transcriptomic analysis, bioinformatic processing and statistical analysis were coordinated by omics2view.consulting GbR, Kiel (Germany).

## Authors contributions

Conceptualization, HS, CP, IKH, KLA, BJMB; Methodology, CP, IKH, HGK, KLA, HS; Data processing, CP, KLA, IKH, MTO, MPH; Investigation, CP, IKH, HGK, KLA; Writing & Review, HS, BJMB, CP and KLA; Funding acquisition, HS, BJMB and MPH; Resources, HS and BJMB.

## Data Availability Statement

The datasets generated during and/or analyzed during the current study are available in Supplementary Tables S1 and S2 and in the XXXX repository.

## Supplementary Files

**Supplementary Information (PDF)**

Material and methods including details of nematode and bacterial strains, mesocosm experiment, common garden experiment, assessment of population growth rate, 16S and ITS amplicon sequencing, microbiome data analysis, RNAseq for transcriptome analysis, and transcriptome data analysis. Supplementary Figures S1-S11 showing results for worm length/area, microbiome composition, and differential gene expression.

**Supplementary Table S1 (MS Excel)**

Supplementary Tables S1-1 to S1-14 with information regarding species included in the CeMbio43 bacterial community, data for population growth rates, worm length and worm area from common garden experiments, male frequencies, test statistics for population growth rates, worm length and worm area from common garden experiments.

**Supplementary Table S2 (MS Excel)**

Supplementary Tables S2-1 to S2-5 including read counts of fungal (ITS2) and bacterial (16S) ASVs from common garden experiment, and results of PerMANOVA tests for differences in microbiomes.

**Supplementary Table S3 (MS Excel)**

Supplementary Tables S3-1 to S3-6 including information on condensed clusters of differentially expressed genes and results of the enrichment analysis with the *C. elegans*-tailored WormExp database.

**Supplementary Movie 1 (QuickTime Movie)**

Video of a sample from the upper compost layer at day 244 of the mesocosm experiment showing large numbers of proliferating worms.

**Supplementary Movie 2 (QuickTime Movie)**

Video of worms in from mesocosm experiment in M9-buffer. The worms are released from a compost sample after the addition of M9-buffer and can thus be secured for further analysis.

**Supplementary Movie 3 (QuickTime Movie)**

Video of worms from mesocosm experiment in M9-buffer. The worms are released from a compost sample after the addition of M9-buffer and can thus be secured for further analysis.

