## Supplementary material for "Host and microbiome jointly contribute to adaptation to a complex environment": Supplemantary Information

[*]Shared first authors

[†]Shared senior authors

Corresponding author:

[1] Department of Evolutionary Ecology and Genetics, Kiel University, Kiel, Germany

[2] Institute of Ecology and Evolution, University of Oregon, Eugene, OR, USA

[3] Institute of Clinical Molecular Biology, Kiel University, Kiel, Germany

[4] Max-Planck Institute for Evolutionary Biology, Ploen, Germany

### Supplementary Materials and Methods

#### Nematode strains

We used the *C. elegans* population $A_0$ (1) which was repeatedly used in various evolution experiments (1–4). Unless otherwise stated, the $A_0$ population was maintained on nematode growth medium (NGM) plates with *Escherichia coli* strain OP50 and synchronized by bleaching following standard procedures (5).

#### Bacterial strains

To prepare the CeMbio43 community, bacteria were thawed from frozen stocks, individual bacteria were cultured in tryptic soy broth (TSB) in deep well microtiter plates for 42 h on a circular shaker at 20 °C, then mixed in equal cell numbers, and the bacterial mixture adjusted to $OD_{600}$10 in PBS.

#### Mesocosm experiment

Lab compost was prepared by placing 100 g autoclaved compost soil in a sterile box. Each box was closed with a lid with three 3 cm holes closed with a sterile foam plug to allow aeration. Potatoes, apples, kohlrabi and carrots including leaves were washed with water, chopped, mixed and approximately 100 g of the plant material was added to each box and mixed with the soil in the box. One ml of freshly prepared CeMbio43 in $OD_{600}$5 was added directly, and another 1 ml and 1.3 ml were added after 24 h and 48 h, respectively. Compost and bacteria were mixed and moistened with 1 to 2 ml of sterile water. Subsequently, approximately 3200 *C. elegans* $A_0$ in different stages were added to the compost. The boxes were stored on a table in a separate room at room temperature. Fresh plant material was added every other week and each compost mixed every week using a sterile wooden spatula. The position of the boxes on the table was shuffled weekly. In order to mimic a natural compost environment, the boxes were not maintained under strictly sterile conditions; therefore, microorganisms (in addition to the initial CeMbio43 inoculum) were regularly introduced to the environment (e.g. via fresh plant material).

After 100 days, we sampled nematodes and bacteria from each box. To collect the nematodes, a randomly chosen subsample of each compost was placed in a 9-cm petri dish, followed by addition of 25 ml of M9-buffer with 0.025% (v/v) of Triton X100 (M9-T). We then transferred 150 µl of worm-containing buffer onto peptone-free NGM (PFM) plates. The resulting worm population was bleached to remove any bacteria, following standard protocols (5), and the surviving eggs were kept overnight in M9-buffer on a shaker at 20 °C and the hatched L1 larvae frozen in a final concentration on 15% (w/v) glycerol in S-buffer at -80 °C. Microbial communities were obtained from the same subsample in M9-buffer by transferring 300 µl of the mixture to a 2 ml microtube containing approximately ten sterile 1 mm zirconia beads, followed by homogenization of the material using a bead ruptor (Bead Ruptor 96, Omni International,

Kennesaw, Georgia, USA) for 3 min at 30 Hz, and conservation of microbes in 10% (v/v) dimethyl sulfoxide (DMSO) at -80 °C.

**Common garden experiment and assessment of nematode population growth rate**

Laboratory compost consisted of approximately 10 g water-washed, grated potatoes, apples, kohlrabi and carrots including greens placed in sterile 60 ml containers. Each laboratory compost was inoculated with 1 ml of CeMbio43 or one of the microbial communities isolated at day 100 from the mesocosms, always standardized to an $OD_{600}$10. The containers were covered with an air-permeable film and then left for three days at room temperature and stirred daily. The NGM agar plates were inoculated with the same CeMbio43 or day-100 microbial communities 24 h before the experiment. At the start of the common garden experiment, we added 100 synchronized *C. elegans* at the fourth larval stage from the ancestral or day-100 host populations in M9-T to the compost or NGM plate replicates. For the second experiment, worm populations were prepared to be devoid of males, thus consisting of hermaphrodites only, in order to ensure comparability of the different *C. elegans* populations. These male-free populations were frozen in aliquots for later usage. They were rechecked for males after thawing and the subsequent initiation of worm cultures with individual L4 nematodes, followed by an assessment of the populations after 5 and 7 days, revealing and confirming the complete absence of males (Supplementary Table S1). Compost and plates were stored at 20 °C and the compost mixed daily by gentle shaking. After five or four days for compost or plates respectively, the worms were collected.

For the analysis of the plates, we collected worms in 5 ml M9-T, centrifuged for 1 min at 500 rpm, the supernatant was removed, and the worms washed four more times in fresh M9-T. The washed worm pellet was split in two microtubes. One microtube was stored at -20 °C until population growth rate, worm length, and worm area were determined. The other tube was stored at -80 °C.

For the compost, nematodes and microbial communities were isolated from a randomly chosen subsample following a standardized protocol. For this, the compost was mixed using a sterile wooden spatula and approximately 0.6 g compost was transferred onto a 9 cm petri dish. 15 ml sterile M9-T were distributed over the sample by gently circling the Petri dish five times. We then collected worms in 3 x 500 µl buffer from a distance of 1 cm, 2 cm and 3 cm around the undissolved compost sample and combined the samples into one microtube. Three technical replicates (three Petri dishes containing a 0.6 g compost subsample) were taken from each compost to compensate for variations in worm counts within a compost sample. Worm samples were frozen in 1.5 ml M9-T at -20 °C for later phenotypic analysis. Population growth rate was determined by first counting worms in 1 µl to 200 µl (depending on worm density) of the frozen sample, repeated a total of three times and calculation of the average. Offspring per worm was calculated by dividing the count result by the 100 worms initially added to each replicate.

The counted worms were extrapolated to the total frozen buffer and, for the compost experiment, to the total compost weight. Worm length and area were determined in Image J (version 2.3.0) using pictures taken with a Leica stereomicroscope (Leica Microsystems GmbH, Wetzlar, Germany). All experiments were performed without knowledge of the origin of the nematodes and bacteria, using codes to avoid observer bias. All treatment combinations were evaluated in parallel and in randomized order.

The population growth rates, worm length and area of different treatment groups were compared with a Wilcoxon rank sum test and Bonferroni correction for multiple comparisons or with an ANOVA. All statistical calculations were performed with R studio software (version 2022.07.2) and can be found in Supplementary Table 1. Graphs were produced with R Studio and edited with Inkscape (version 1.1).

**16S and ITS amplicon sequencing for microbiome analysis of common garden experiment**

The compost was mixed with a sterile, wooden spatula and a compost (i.e., substrate) sample was collected and frozen at -20 °C until further use. Worms were collected from the compost sample following a previously published protocol (6). Briefly, a compost sample was covered with M9-T and emerging worms were collected using a pipette. The worms were transferred to sterile M9-T in a 3 cm Petri dish and then to a sterile microtube and kept for at least 2 min in 10 mM Tetramisole to stop ingestion and excretion of bacteria. Worms were washed another four times in fresh M9-T and frozen in 300 µl of M9-T in a 2-ml microtube in liquid nitrogen and finally stored at -80 °C.

For isolation of bacterial DNA from nematodes, we added five to ten sterile 1 mm zirconia beads to the thawed worm samples. After crushing the samples for 3 minutes at 30 Hz, each sample was transferred to a 1.5 ml microtube and centrifuged at 8000 rpm for 3 minutes. All but 100 µl of the supernatant was removed and the pellet was resuspended in the remaining 100 µl. Using a tissue kit (Macherey-Nagel, Düren, Germany), DNA was isolated according to the manufacturer's instructions and stored at -20 °C.

For DNA isolation from compost, we shredded approx. 100 µl of a compost sample with 5-10 sterile 1 mm zirconia beads and 300 µl RNase-free water in a 2 ml microtube for 3 min at 30 Hz. The tubes were briefly centrifuged to spin down larger substrate particles. DNA was isolated from 100 µl compost supernatant using a modified Cetyl Trimethyl Ammonium Bromide- (CTAB) based protocol (7,8).

16S rRNA gene amplicon libraries of worm and compost DNA samples were prepared using the primers 341F (5’CCTACGGGNGGCWGCAG-3’) and 806R (5’GACTACHVGGGTATCTAATCC-3’) covering the V3–V4 region of the 16S rRNA gene. Libraries were sequenced on the Miseq platform using the v3-Chemie 2 × 300 bp. ITS2 libraries were prepared using the primers 5.8S-Fun (5’AACTTTYRRCAAYGGATCWCT-3’) and ITS4-Fun (5’AGCCTCCGCTTATTGATATGCTTAART-3’) (9). The ITS2 libraries were sequenced on the Miseq platform using the v2 nano kit 2 x 250 bp. The microbial community composition was analyzed for four replicates.

**Microbiome data analysis**

Raw amplicon sequencing reads were processed using the QIIME 2 v2022.2 microbiome bioinformatics platform (10). Primer and adapter sequences were removed with cutadapt (11). For the 16S amplicons, reads were filtered based on quality scores and forward and reverse reads joined with vsearch (12), and amplicon sequence variants (ASVs) were resolved with deblur (13). Taxonomy was assigned to the 16S ASVs with a naïve Bayes classifier pre-trained on the Silva 138 SSU database (14–16). For the ITS amplicons, ASVs were resolved from the forward reads with the DADA2 pipeline which removes low quality and chimeric sequences (17). Taxonomy was assigned to the ITS ASVs with a naïve Bayes classifier pre-trained on the UNITE v9.0 database (14,15,18,19). The raw reads are available from the Sequence Read Archive (SRA) database under accession number XXXX.

All statistical analyses of the microbiome data were conducted in R Studio (20,21). Potential contaminant ASVs for both the 16S and ITS datasets were identified with the *decontam* package using the prevalence method and removed (22). We then assessed differences in microbial community composition in three main sets of analyses, which addressed the following questions:

(i) How do Box 1 and Box 2 substrate microbiomes differ in taxonomic composition, and to what degree are substrate microbiome members selected by worms? To address this question, we compared microbiome composition among substrates to which ancestral worms and either the reference microbiome (including the CeMbio43 bacterial community), Box 1 day-100 microbiome or Box 2 day-100 microbiome were added and used differences in microbiome composition between these substrates and their respective worms as an indication of selection.

(ii) Which microbiome members are associated with the increase in fitness observed for Box 1 worms colonized by the corresponding Box 1 microbiome? To address this question, we compared microbiomes between ancestral and Box 1 day-100 worms exposed to the Box 1 day-100 microbiome and compared these worms to their respective substrates.

(iii) Which members of the Box 2 microbiome are associated with low worm fitness? Here we used differences between the substrate and worm microbiomes for both Box 2 day-100 and ancestral worms exposed to the Box 2 day-100 microbiome as an indication of selection by the worm and compared the selected microbiomes of Box 2 day-100 and ancestral worms exposed to the Box 2 day-100 microbiomes.

To conduct these analyses, we first quantified dissimilarity in microbiome composition between samples with robust Aitchison distance (23) and visualized these relationships with ordinations of principal coordinates analyses. We partitioned variation in Aitchison distance among worm type, microbiome type, sample type and their interactions and tested for significance with permutational multivariate analysis of variance (PerMANOVA) (24,25). To determine which amplicon sequence variants differed in relative abundance between pairs of treatment combinations, we used the ALDex2 R package (v 1.31.0) and considered ASVs with an effect size > 1 or < -1 to be differently abundant (26–28).

**RNAseq for transcriptome analysis of *C. elegans* populations**

Approximately 1000 worms were added to the compost. After 24 h, a compost subsample was placed in a 9-cm Petri dish and covered with M9-T. The emerging worms were transferred three times to fresh M9-T in a 3 cm Petri dish in as little liquid as possible to remove bacteria adhering to the worms. At least 50 worms were frozen in 800 µl TRIzol Reagent (Thermo Fisher Scientific, Waltham, MA, United States) in a 2-ml microtube in liquid nitrogen. Worms in TRIzol were thawed five times at 45 °C and refrozen in liquid nitrogen to break up the worm cuticle. Total RNA was isolated using a Direct-zol RNA MicroPrep Kit (Zymo Research, Irvine, CA, United States) following manufacturer's instructions and stored at -80 °C. The transcriptome was analyzed for five replicates. RNA libraries were prepared for sequencing using the Illumina stranded total RNA kit with RiboZero Plus (Illumina, San Diego, USA, catalog number 20040529) according to the manufacturer's protocol. Libraries were sequenced on an Illumina NovaSeq 6000 with paired-end strategy and read length of 100 bp (Illumina, San Diego, USA, NovaSeq 6000 SP Reagent Kit v1.5 (200 cycles), catalog number 20040719). The raw data is available from the GEO database (Barrett et al., 2012; Edgar et al., 2002) under the GSE number XXXX.

**Transcriptome data analysis**

The obtained sequence data was first processed, checked for quality, and filtered. In detail, we removed ribosomal and transfer RNA sequences and, thereafter, contaminant reads with the program BBSplit of the BBTools package v38.45 (29). Read quality trimming was performed with the BBTools package v38.45 (29), including removal of duplicate reads, adapter sequences, low-entropy reads, and trimming of bases with quality scores < 10. Reads with invalid or ambiguous bases and reads with a length < 50 base pairs (bp) were discarded. Only reads surviving quality trimming as pairs entered downstream analysis. Read quality recalibration and error correction was performed with the BBTools package v38.45 (29). The filtered reads were mapped with STAR v2.7.10b (30) to the reference genome of *C. elegans* strain N2, release WS286, retrieved from WormBase (31). We then assessed differential gene expression in three main sets of analyses, which addressed the following three main questions:

(i) To what extent do the Box 1 day-100 and Box 2 day-100 populations differ from each and the ancestral *C. elegans* population and thus indicate genetic evolution? To address this question, we compared variation in gene expression among the three worm populations upon exposure to the same reference microbiomes (including the CeMbio43 bacterial community) in compost, thus ensuring identical growth conditions for the three populations.

(ii) Which gene expression changes characterize the Box 1 day-100 worms colonized by their coexisting Box 1-day 100 microbiome and thus underlie the high fitness expressed by this assemblage in the Box 1 common garden experiment? For this, we identified and characterized the gene expression signature that is unique to the Box 1 day-100 worm - Box 1 day-100 microbiome combination in comparison to the remaining three host-microbiome treatments of the Box 1 common garden experiment.

(iii) Which gene expression changes characterize the low fitness of worms exposed to the Box 2 day-100 microbiome? To address this question, we compared gene expression changes in the two nematode populations exposed to the Box 2 day-100 microbiome to those exposed to the reference microbiome.

All three sets of analyses included the following steps. At the beginning, we explored variation in gene expression across the considered treatments using a principal component analysis (PCA) with R package ggplot2 v3.4.0 (32) in R v4.2.1 (20). Thereafter, we inferred differential gene expression, always relative to the combination of the ancestral *C. elegans* population with the reference microbiome, using R package DESeq2 v1.36.0 (33). We only included genes in the analysis if they (i) have an existing Entrez gene symbol, (ii) are coding, (iii) are functional (i.e., not a pseudogene), and (iv) have ≥ 10 counts in ≥ 5 samples. Gene count distribution was modeled using a negative binomial generalized linear model. Statistical significance of the treatment effect on individual genes was determined with the Wald test (34). Effect size is expressed as log2(fold-change) of size-factor-normalized gene counts. Probabilities were adjusted using the false discovery rate (35). For the sake of clarity, only genes with a false discovery rate ≤ 0.1 are shown. Genes with absolute log2(fold-change) ≥ 1 and FDR ≤ 0.05 or 0.01, respectively, in at least one contrast, were subjected to *k*-means clustering. The optimal number of clusters was determined based on Akaike's Information Criterion. The process was repeated 100 times to account for random effects, and the median optimal number of clusters was taken as the final result. Gene clusters of contrasts with FDR ≤ 0.01 were further condensed by performing a second round of *k*-means clustering using the cluster medians of the first round as input. The C Index (36) was used to determine the optimal number of second-round clusters. We used the results of the second round of *k*-means clustering for subsequent enrichment analyses.

For enrichment analyses, we focused on individual gene clusters that showed the respective transcriptome signatures needed to address the three above questions (e.g., gene clusters indicating a difference in transcriptome response between the Box 1 and Box 2 worm populations to address question (i)). Two types of enrichment analyses were performed. On the one hand, we used the *Database for Annotation, Visualization and Integrated Discovery* (DAVID; (37)) and studied enrichment according to Gene Ontology (GO) (38), including only categories with a probability ≤ 0.05 (after FDR correction). Cluster importance of a category was defined as Importance, following:

$$Importance_{ij} = 100 \times \frac{H_{ij}}{N_i} \times \frac{N_i}{N}$$

where $H_{ij}$ is the number of genes in cluster *i* with a hit in GO category *j*, $N_i$ is the total number of genes in cluster *i*, and *N* is the overall number of genes in all clusters. The rationale of this approach is to normalize the proportion of hits within each cluster by relative cluster size to avoid bias due to the latter. Cluster importance of GO categories is visualized as heatmaps.

On the other hand, we performed an enrichment with the *C. elegans*-specific gene expression database WormExp (39), which contains approx. 3000 published gene expression data sets for *C. elegans* under diverse conditions, allowing a more taxon-specific inquiry of enriched expression categories. Probabilities of enriched gene sets were adjusted using FDR, and the relationship of enriched gene sets was evaluated using hierarchical clustering, visualized via heatmaps.

**Supplementary Figures**

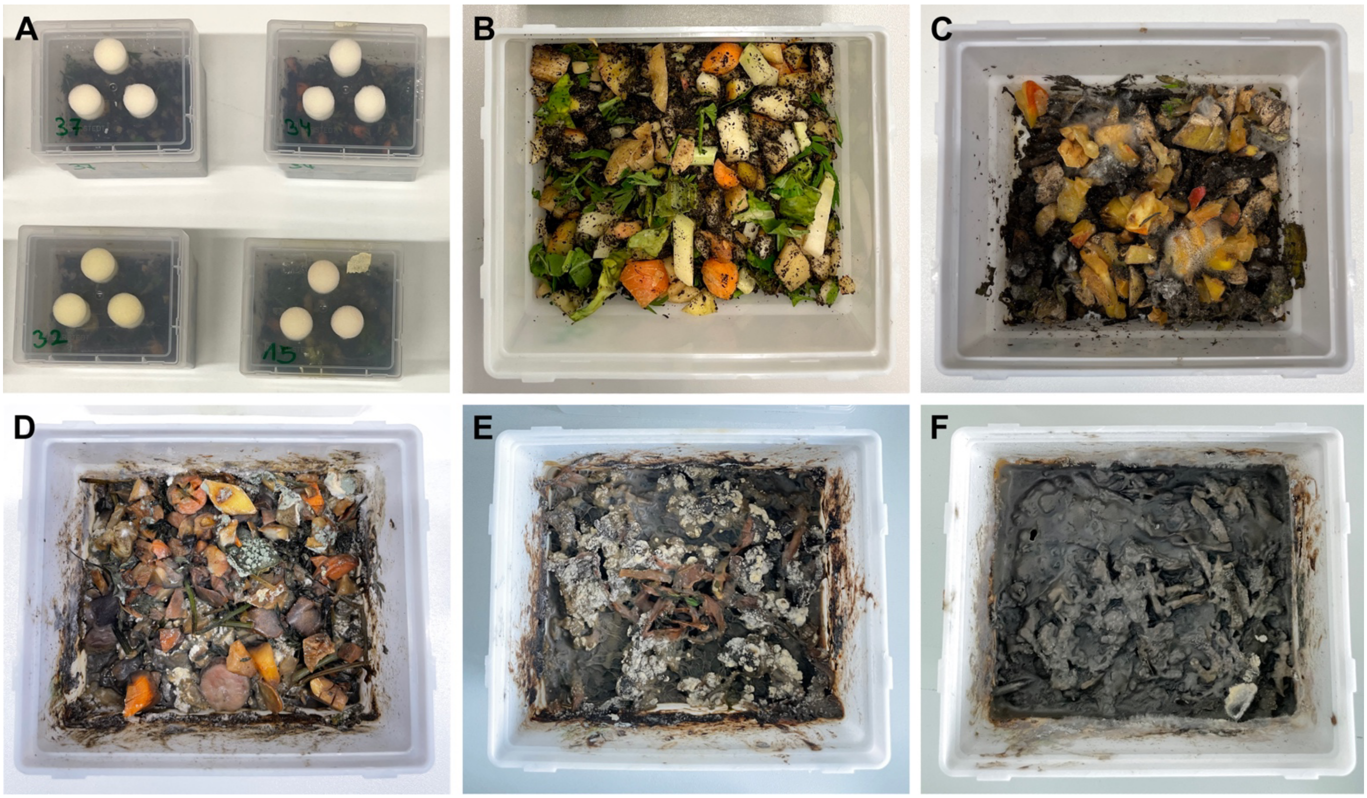


**Figure S1: Mesocosm experiment in laboratory compost. (A)** A mesocosm experiment was performed in boxes containing laboratory compost. **(B)** Fresh produce and compost soil were added to the boxes at the beginning of the experiment, followed by the addition of 43 native microbiota bacteria (CeMbio43) and a genetically diverse worm population. The laboratory compost was supplemented with fresh produce every two weeks. **(C)** After two weeks and **(D)** ten weeks, new microbes were visible and the compost showed signs of decomposition. On day 100, worms and microbes were harvested from **(E)** Box 1 and **(F)** Box 2.

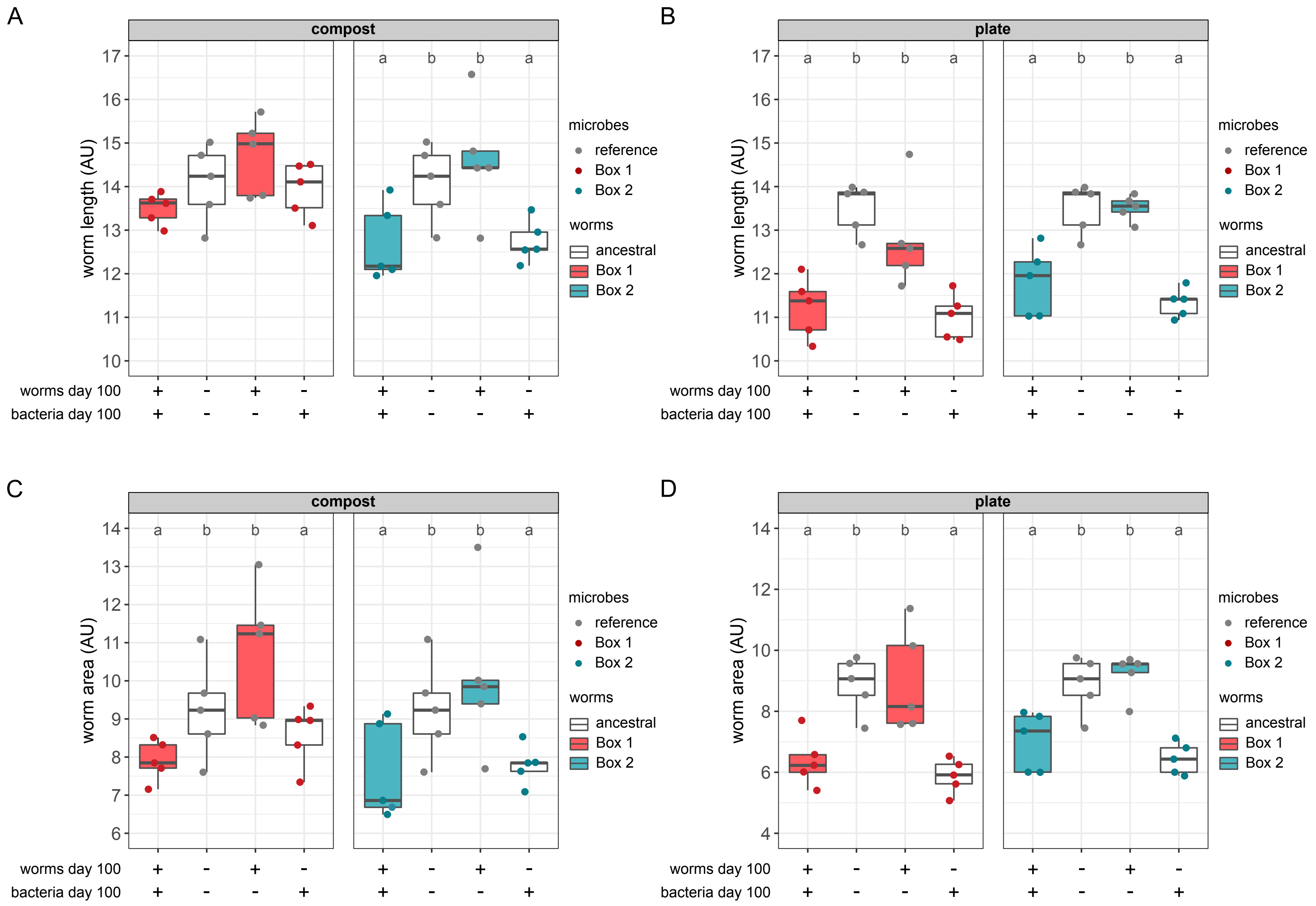


**Figure S2: Host and microbiome can jointly determine nematode fitness in the novel compost environment.** Results of common garden experiments, in which worm length and worm area was measured for *C. elegans* populations isolated from mesocosms at day 100 (+) and ancestral worms (-) in the presence of mesocosm day-100 microbiomes (+) or reference microbiomes including the CeMbio43 bacterial community (-). Worm length was measured for worms from **(A)** compost and **(B)** plates. Worm area was measured for worms from **(C)** compost and **(D)** plates. Worm length and worm area are shown in arbitrary units (AU) for Box 1 (red boxes), Box 2 (blue boxes), and ancestral worm populations (white boxes) in the presence of Box 1 (red dots) or Box 2 (blue dots) microbiomes or reference microbiomes (gray dots). Results are summarized as boxplots with the median as a thick horizontal line, the interquartile range as box, the whiskers as vertical lines, and each replicate depicted by a dot or symbol. n = 5.

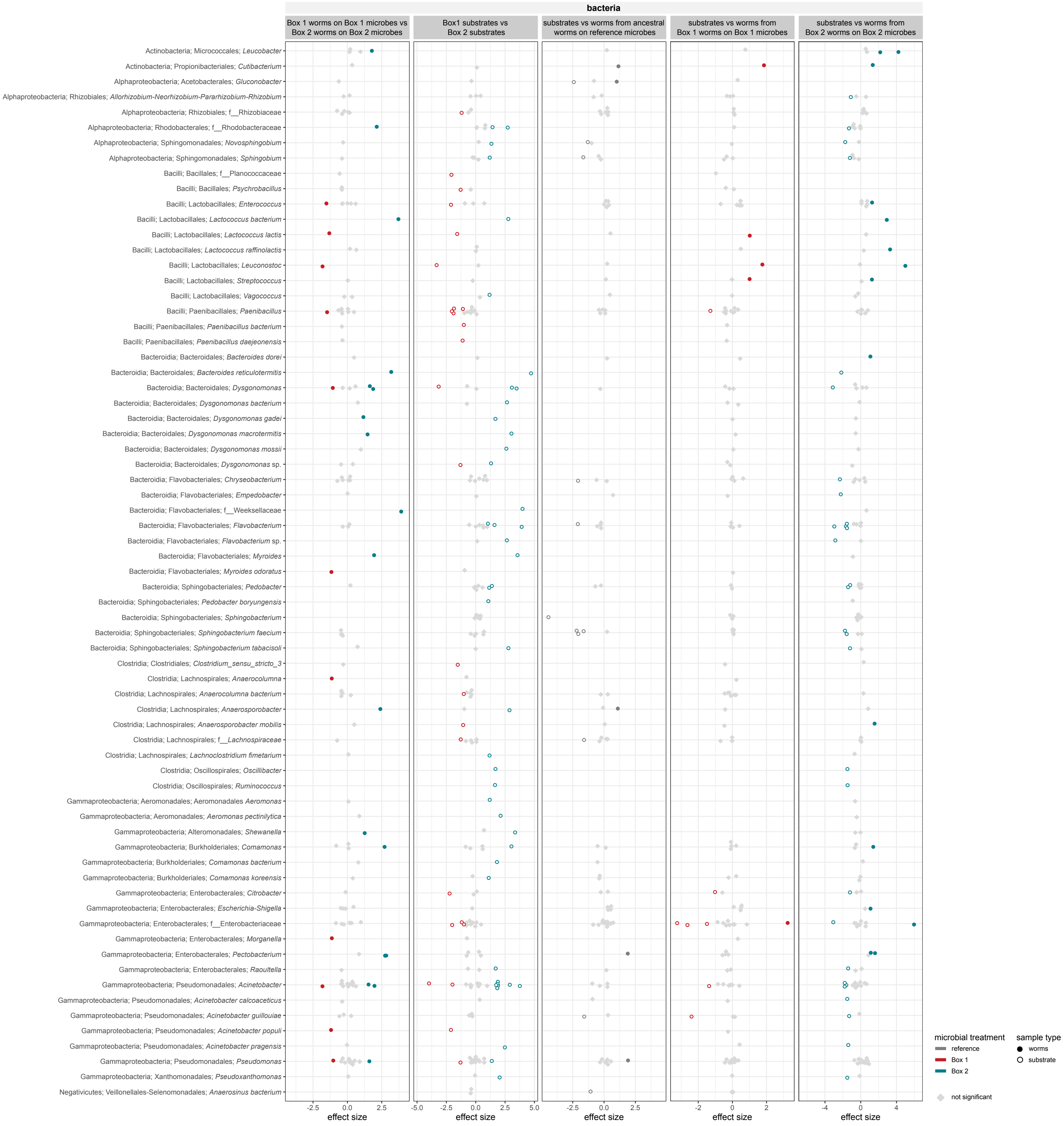


**Figure S3: Microbiome treatments resulted in differences in both compost and nematode microbiomes**. Differential abundance analysis of worm/substrate bacterial microbiomes (Box 1 day-100, Box 2 day-100 or references inoculum including the CeMbio43 bacterial community.

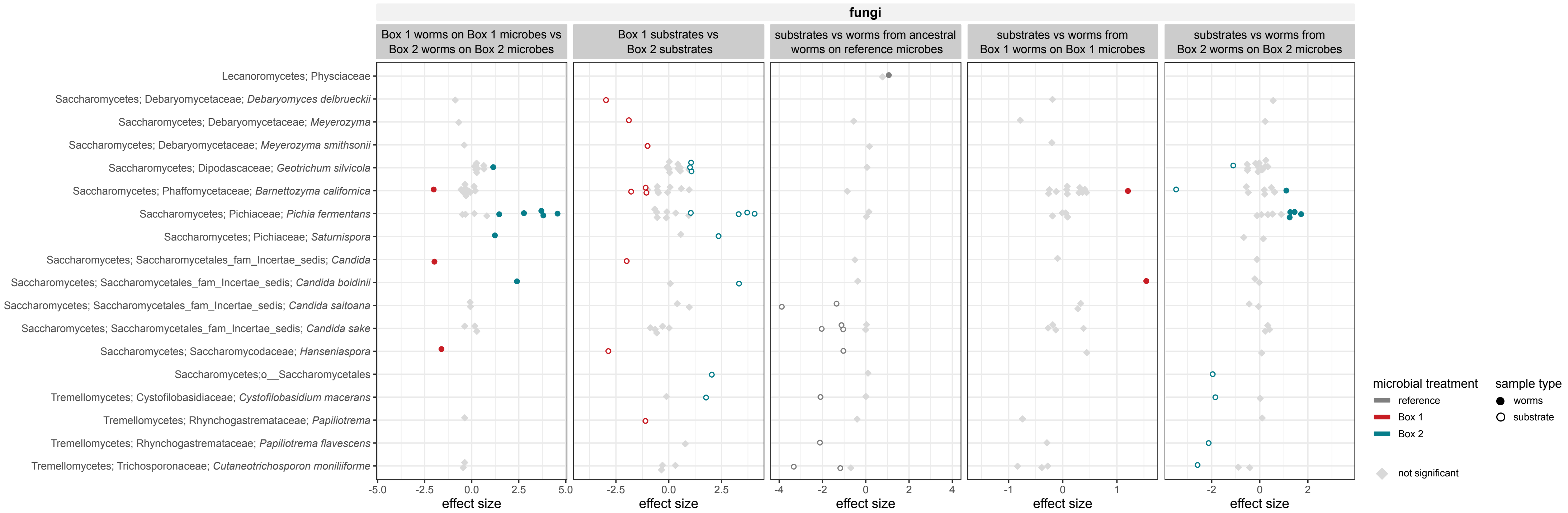


**Figure S4: Microbiome treatments resulted in differences in both compost and nematode microbiomes**. Differential abundance analysis of worm/substrate fungal microbiomes (Box 1 day-100, Box 2 day-100 or reference inoculum including the CeMbio43 bacterial community).

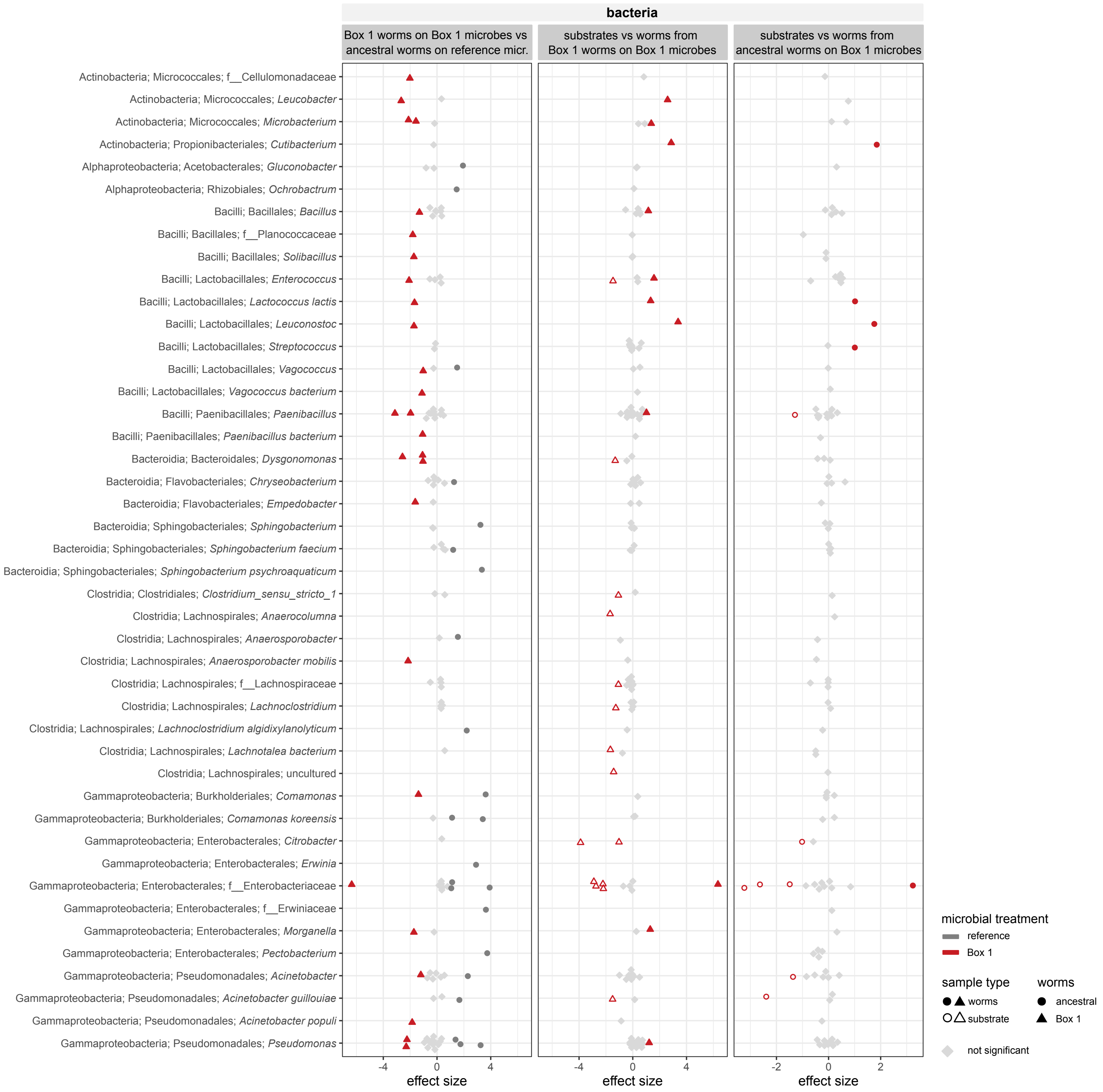


**Figure S5: Differences in microbiome composition were associated with increased fitness in nematodes from the Box 1 mesocosm.** Differential abundance analysis of bacterial microbiomes from ancestral or Box 1 day-100 worms/substrates exposed to Box 1 day-100 inoculum or references inoculum including the CeMbio43 bacterial community.

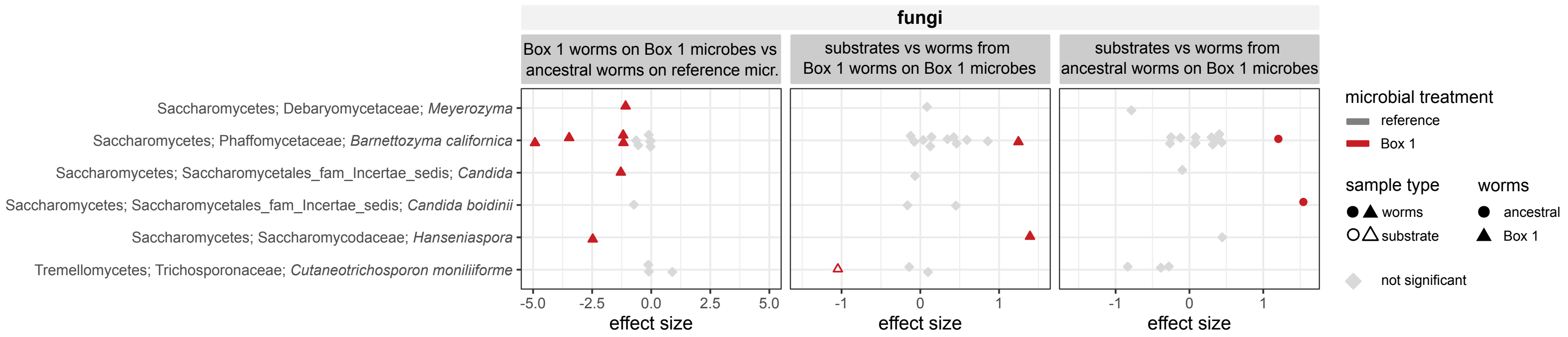


**Figure S6: Differences in microbiome composition were associated with increased fitness in nematodes from the Box 1 mesocosm.** Differential abundance analysis of fungal microbiomes from ancestral or Box 1 day-100 worms/substrates exposed to Box 1 day-100 inoculum or reference inoculum including the CeMbio43 bacterial community.

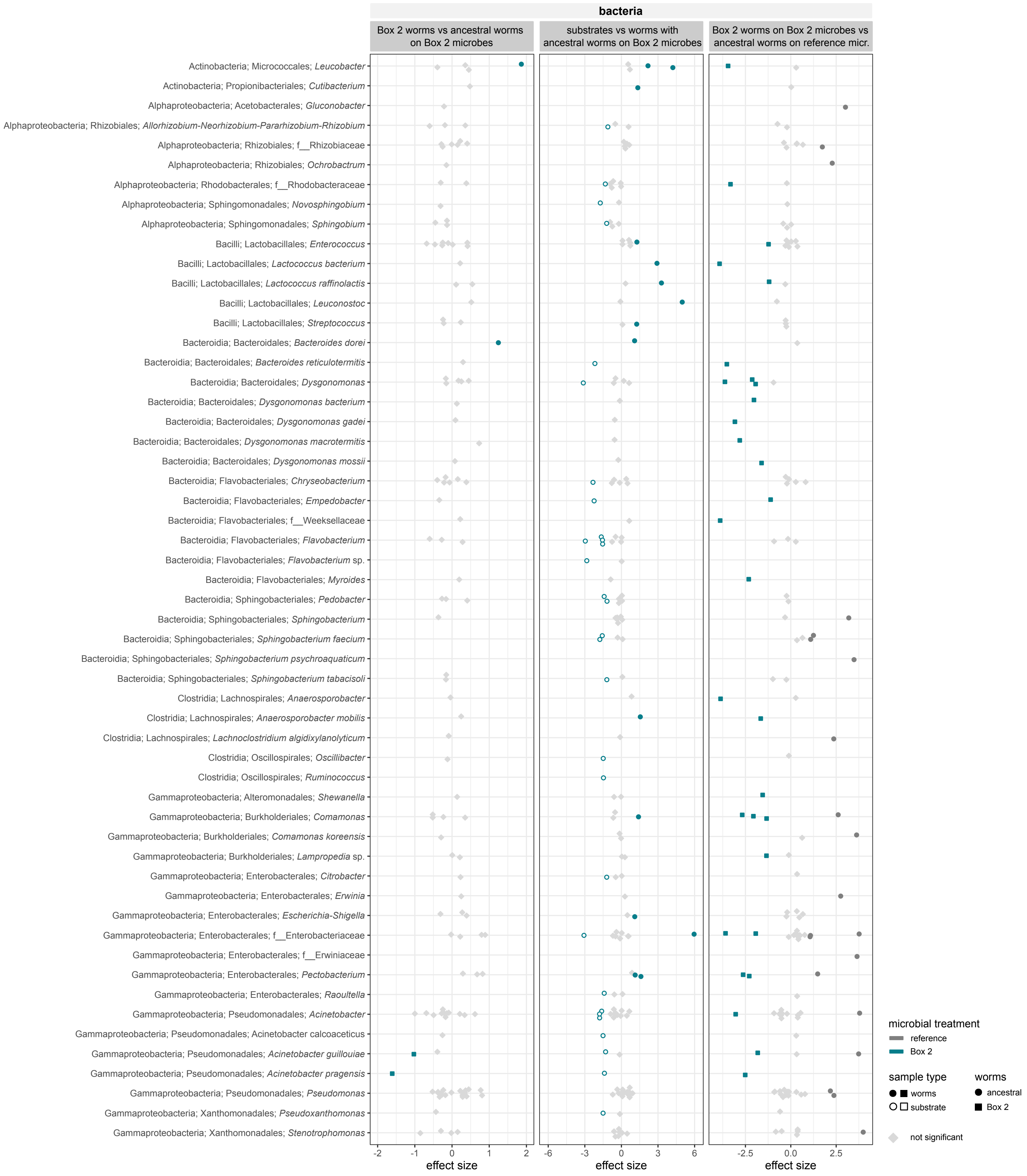


**Figure S7: Differences in microbiome composition were associated with decreased fitness in nematodes from the Box 2 mesocosm.** Differential abundance analysis of bacterial microbiomes from ancestral or Box 2 day-100 worms/substrates exposed to Box 2 day-100 inoculum or reference inoculum including the CeMbio43 bacterial community.

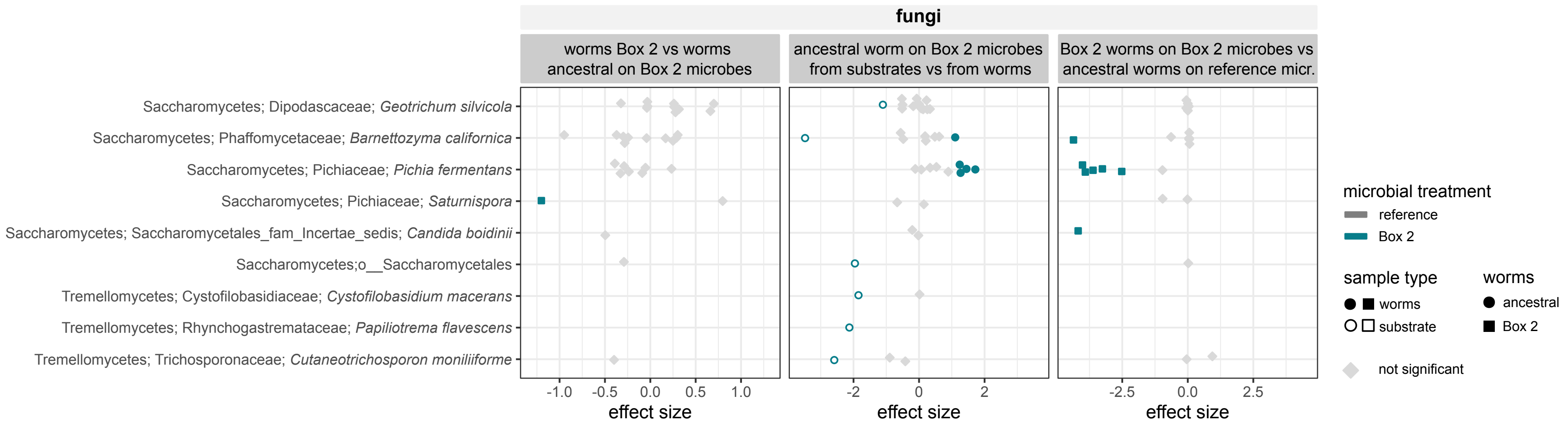


**Figure S8: Differences in microbiome composition were associated with decreased fitness in nematodes from the Box 2 mesocosm.** Differential abundance analysis of fungal microbiomes from ancestral or Box 2 day-100 worms/substrates exposed to Box 2 day-100 inoculum or reference inoculum.

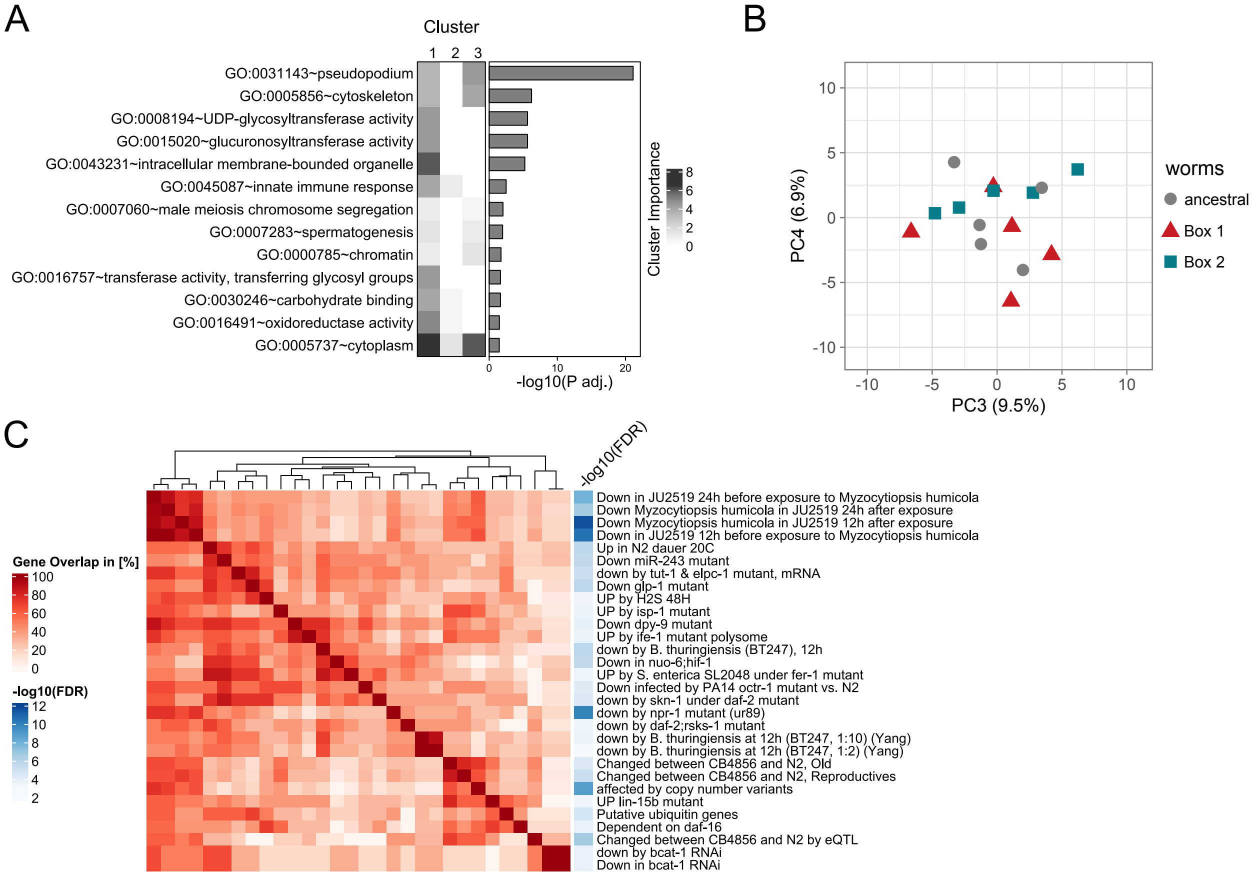


**Figure S9: Differential gene expression in the adapted Box 1 *C. elegans* populations.** Transcriptome data analysis for the comparison of ancestral, Box 1 day-100 and Box 2 day-100 *C. elegans* populations assayed under identical compost conditions with reference microbiomes including the CeMbio43 bacterial community. **(A)** Enriched gene ontology (GO) terms of differentially expressed genes. GO enrichment analysis was performed by DAVID. **(B)** General variation in gene expression was explored with a principal component analysis, whereby the panel shows the spread of sample variation along the third and fourth principal components (PC3, PC4). **(C)** shows the results of the focused enrichment analysis of cluster 2 with the *C. elegans*-tailored WormExp database and visualization of differential expression using heatmaps, whereby the heatmaps always show the gene overlap in percent. Description on the right gives terms of the gene functions and fold change after FDR correction.

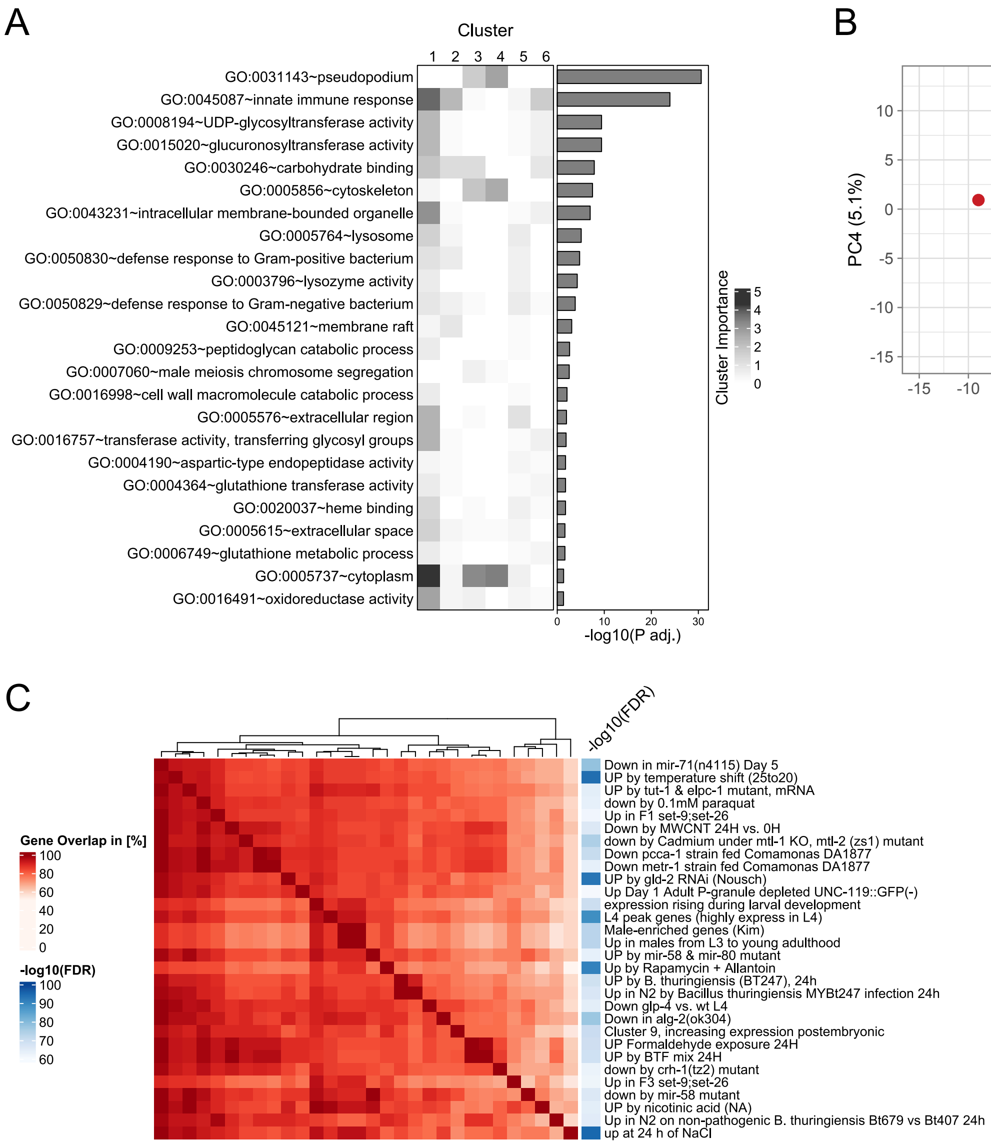


**Figure S10: Differential gene expression in the adapted Box 1 *C. elegans* populations.** Transcriptome data analysis for the comparison of all possible host-microbiome combinations for the Box 1 common garden experiment. Ancestral or Box 1 worms were combined with either the reference microbes (including the CeMbio43 bacterial community) or the Box 1 day-100 microbes. **(A)** Enriched gene ontology (GO) terms of differentially expressed genes. GO enrichment analysis was performed by DAVID. **(B)** General variation in gene expression was explored with a principal component analysis, whereby the panel shows the spread of sample variation along the third and fourth principal components (PC3, PC4). **(C)** shows the results of the focused enrichment analysis of cluster 3 with the *C. elegans*-tailored WormExp database and visualization of differential expression using heatmaps, whereby the heatmaps always show the gene overlap in percent. Description on the right gives terms of the gene functions and fold change after FDR correction.

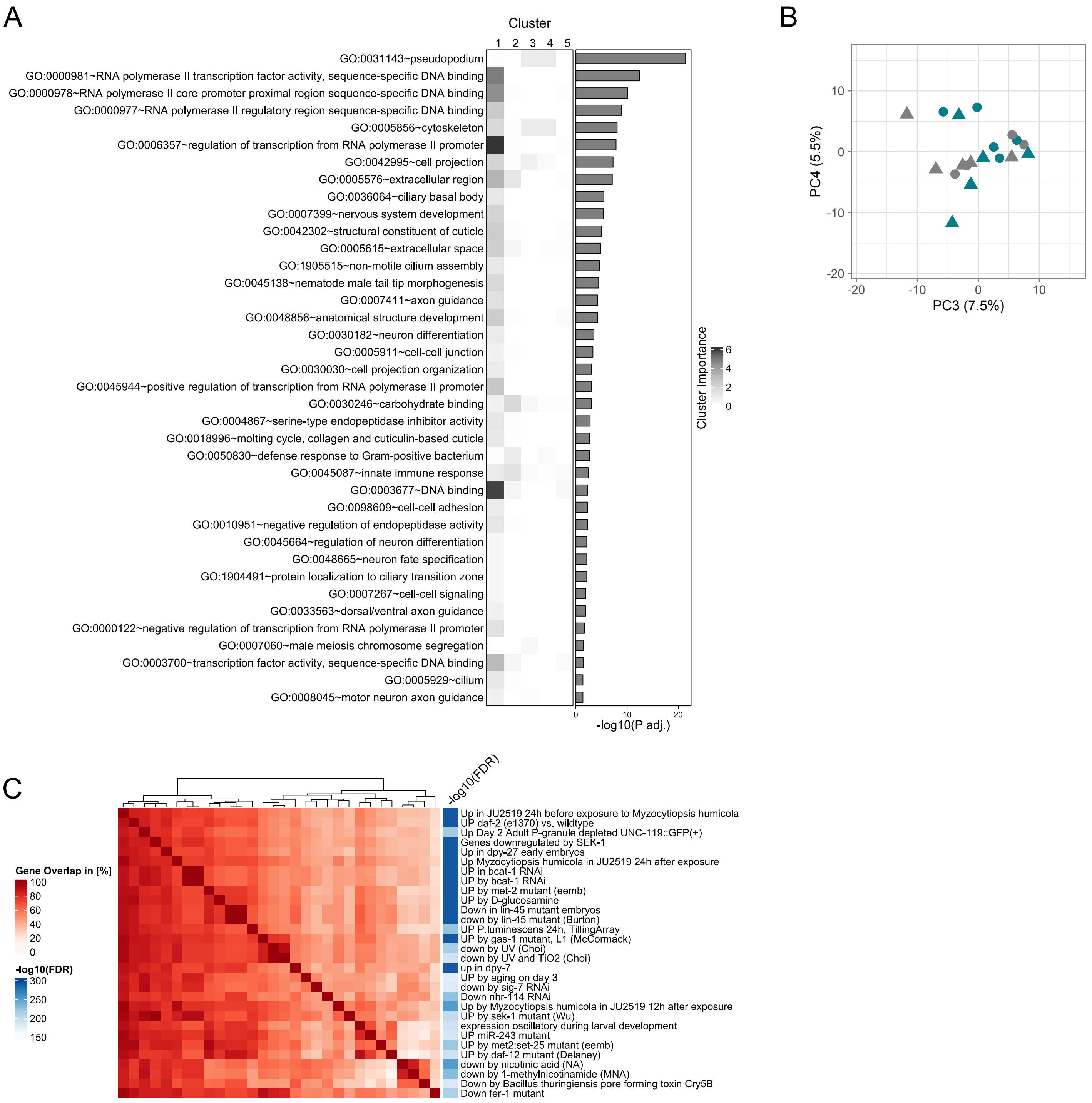


**Figure S11: Differential gene expression in the adapted Box 2 *C. elegans* populations.** Transcriptome data analysis for the comparison of all possible host-microbiome combinations for the Box 2 common garden experiment. Ancestral or Box 2 day-100 worms were combined with either the reference microbes (including the CeMbio43 bacterial community) or the Box 2 day-100 microbes. **(A)** Enriched gene ontology (GO) terms of differentially expressed genes. GO enrichment analysis was performed by DAVID. **(B)** General variation in gene expression was explored with a principal component analysis, whereby the panel shows the spread of sample variation along the third and fourth principal components (PC3, PC4). **(C)** shows the results of the focused enrichment analysis of cluster 1 with the *C. elegans*-tailored WormExp database and visualization of differential expression using heatmaps, whereby the heatmaps always show the gene overlap in percent. Description on the right gives terms of the gene functions and fold change after FDR correction.
